# Tetrapod diversity facets in jeopardy during the Anthropocene

**DOI:** 10.1101/2021.07.01.450689

**Authors:** Enrico Tordoni, Aurèle Toussaint, Meelis Pärtel, David Nogues-Bravo, Spyros Theodoridis, Carlos Pérez Carmona

## Abstract

Human activities have eroded biodiversity, yet the varying influence of past versus recent impacts across the distinct facets of biodiversity is still poorly understood. Weighting taxonomic information by phylogenetic and functional diversity in a novel multifaceted index (μ-Diversity) across more than 17,000 tetrapod species, we show the geography of multifaceted tetrapod diversity, and the role of climate stability and water-energy dynamics coupled with the timing of inception of agriculture in explaining broad-scale patterns of tetrapod diversity. In particular, the varying geography of the timing of agriculture expansion since the Neolithic affected μ-Diversity at least as much as recent human impacts, especially in birds, mammals, and reptiles, suggesting that human imprints may have shaped tetrapod diversity for millennia through legacy effects of past land use modifications. The long-lasting effect of humans will only accelerate, as the most diverse areas for μ-Diversity (tropical Africa, South East Asia and Central and South America) are disproportionally exposed to both future climate and land-use change.

## Main text

As human appropriation of natural ecosystems and climate change accelerate, the risk of losing species and their associated functions increases and jeopardizes the adaptive capacity of ecological systems worldwide. A recent assessment^1^ indicates that 77% of lands have been already modified by humans; but the interaction between climate and land use change will likely promote large disruptions in natural ecosystems in the near future^2–4^. To efficiently protect biodiversity from these anthropogenic threats and to anticipate the future trajectories of global change, we need to gain a better understanding of the drivers that shape its global geography across all biodiversity levels and taxa. Until the last decade, studies seeking to understand broad-scale diversity patterns mostly focused on the number of species alone (taxonomic diversity – TD) of individual taxonomic groups, but biodiversity is a multidimensional metric encompassing multiple dimensions. In this sense, a comprehensive evaluation requires considering other diversity facets such as evolutionary history (phylogenetic diversity ̶ PD^5^) and trait differences among species (functional diversity ̶ FD^6^) and possibly their integration, since we still lack a synergistic estimation of the overall biodiversity along with its main drivers. This evaluation would be beneficial for biodiversity conservation^7^ and may avoid the selective prioritization of specific targets for conservation (e.g., endemic species, vulnerability, PD)^8, 9^.

Water-energy dynamics and climate/biome stability seem to play a prominent role to explain broad-scale patterns of life^10, 11^, influencing population fluctuations, speciation and extinction events. While water-energy dynamics predict higher richness in warmer and more humid environments^12, 13^ (i.e., the tropics), the stability of climate since at least Late Quaternary seems to have played an important role in shaping species distributions. Indeed, long-term climatic stability may have been crucial releasing species from disturbances (e.g., climatic oscillations from wet to dry periods), and promoting the accumulation of older lineages and a longer persistence of biomes (i.e., niche conservatism’^14^), which should be reflected by a higher PD and an overall higher species diversity^15, 16^. Likewise, higher FD is also expected in warmer and productive environments where water availability is not a limiting factor^17^, since some traits combinations are not viable in more extreme conditions and niche conservatism may further constrain the set of realized functional strategies outside of their ancestral environments. Finally, humans induced extensive land-use changes in the last decades which have greatly affected global biodiversity^18^, and the exposure of species to global change (i.e., the magnitude and rate of environmental change experienced by a species in a given location^19^) has steadily increased since 1970s, further exacerbating the global biodiversity crisis. Recent research exploiting updated simulations, archeological data and fossil pollen have shown an early human role in triggering land-use changes and modifying vegetation patterns^20, 21^, suggesting that anthropogenic impacts on biodiversity might have deeper roots in time^22–24^. However, it is still unclear up to what point past land-use legacies have influenced different aspects of tetrapod diversity globally, being this information of paramount importance to foresee and mitigate future human impacts. In this sense, there is even far less information on where human impacts will primarily affect all three taxonomic, phylogenetic and functional tetrapod diversity in the near future, and how these aspects of tetrapod diversity might respond to climate and land-use change.

By leveraging recent advances in data availability and computational power, we provide the first integrated analysis of global variation of taxonomic, functional and phylogenetic diversity for almost half of the known extant vertebrate species (17,341 species encompassing 3,912 terrestrial mammals, 3,239 amphibians, 3,338 reptiles and 6,852 birds; Table S1). We develop a new metric (μ-Diversity) weighting taxonomic diversity in a given location by its average phylogenetic and functional diversity (ecological distinctiveness – ED) to obtain a single quantifiable measure that reflects the effective diversity present in an area and can provide a complementary perspective to define prioritization strategy. In a nutshell, our analyses reveal key findings about 1) the geography of multifaceted tetrapod diversity, 2) the primary role of climate stability and energy availability to explain broad-scale patterns μ-Diversity, but we also detected that the timing of inception of agriculture, which was strongly associated to reductions in the ecological differentiation among species, equal or exceed in importance current land use intensity; 3) the exposure of μ-Diversity to recent developments of global projections of future climate and land-use change, showing that the areas of global importance for μ-Diversity are disproportionally exposed to mid-21^st^ century global change.

## Results and discussion

### Global geography of tetrapod μ-Diversity

We estimated tetrapod μ-Diversity, which weights species richness according to the amount of non-redundant functional and evolutionary information present in a given assemblage, thus reflecting a balanced combination of taxonomic, functional and phylogenetic diversity within and across taxonomic groups. Global tetrapod μ-Diversity is highest in tropical sub-Saharan Africa, Central and South-America, South-East Asia and Eastern Australia (Fig. 1). The relatively low correlation between phylogenetic and functional diversity across taxonomic groups (range = 0.01–0.39, Table S2, Fig. S1) implies a spatial mismatch in the global spatial diversity patterns, suggesting also that phylogenetic diversity captures only a portion of functional diversity, in agreement with recent works^25, 26^. South America is extremely rich in taxonomic diversity, but it shows a relatively low ecological distinctiveness, suggesting that these assemblages are characterized by closely related species that tend to share similar set of traits in agreement with ‘phylogenetic niche conservatism’^27^.

**Fig. 1.**
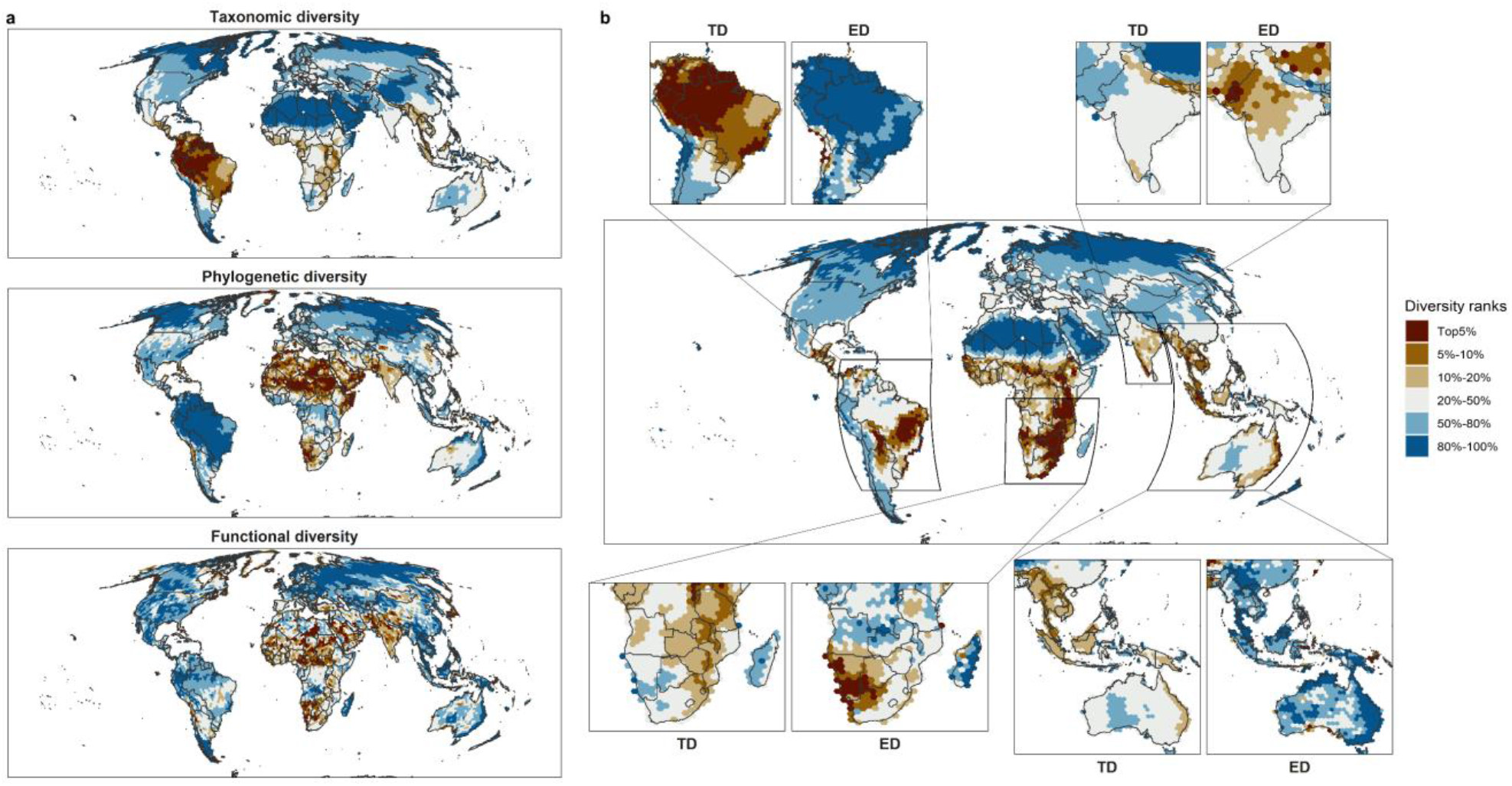
Global areas of importance for tetrapod diversity. **a**, Global pattern of taxonomic, phylogenetic and functional diversity. Tetrapod taxonomic diversity was calculated as the total species richness, while phylogenetic and functional diversity as the median across taxonomic groups. **b**, Global μ-Diversity patterns; insets represent areas of high μ-Diversity where the independent contribution of taxonomic diversity (TD) and ecological distinctiveness (ED) is highlighted. μ-Diversity was expressed weighting species richness by its phylogenetic (sesPD) and functional diversity (sesFD) in each taxonomic group, after having scaled them between 0–1. These were later summed across taxonomic groups to obtain tetrapod μ-Diversity. All diversities were ranked by the most (1–5) to least (80–100) diverse areas globally. Dark brown tones denote the most diverse grid cells while blue tones indicate cold spots.

Despite the relatively low number of tetrapod species with respect to the Neotropics, the Afrotropical and Indo-Malayan realms stand out owing to their higher functional and phylogenetic diversity (Fig. S2). Sub-Saharan tropical Africa and Brazil (e.g., Cerrado and Atlantic forest) contain the highest avian μ-Diversity (Fig. S3a,e,i), followed by eastern Australia and by the Hindukush-Himalaya range^28^. Mammal μ-Diversity (Fig. S3b,f,j) is higher in sub-Saharan tropical Africa, Central America and South-East Asia, whereas high levels of amphibian μ-Diversity (Fig. S3d,h,l) are mainly clustered in Amazonia in agreement with previous studies on these taxonomic groups^29, 30^. Reptiles (Fig. S3c,g,k) displayed the highest diversity in Brazil, Australia, Morocco, South-East Asia and in the Indian subcontinent, highlighting also the crucial role of arid and semi-arid habitats for the their conservation^31^. Indeed, our multi-faceted evaluation reveals that some areas which are usually perceived to support low taxonomic diversity such as arid and semiarid regions (e.g., Sahara-Sahel area, Indian subcontinent) host species with high levels of ecological distinctiveness, especially for birds, mammals, and reptiles (Fig. S3)^32^.

### Energy availability, climate stability, and timing of agriculture expansion affect μ-Diversity

We used random forest models to explore how climate/biome stability, environmental (Potential Evapotranspiration – PET, forest cover, and soil moisture) and past and present anthropogenic factors (year of onset of agriculture encompassing all forms of continuous cultivation^20^ and Human Footprint Index-HFP^33^) have affected the current μ-Diversity patterns considering all tetrapods together and for individual taxonomic groups and diversity facets (i.e. TD and ED). These variables were highly predictive of global patterns in μ-Diversity for all tetrapods combined (R^2^ = 0.80 ± 0.03, Normalized Root Mean Square Error – NRMSE = 0.14 ± 0.03; median ± SD) and, to a lower level, also for individual groups (range NRMSE = 0.14 – 0.19; range R^2^ = 0.69 – 0.76; Table S3). Broadly speaking, μ-Diversity patterns tend to mirror TD in some cases, whereas in other they reflect a trade-off between TD and ED with some differences that are taxa-dependent.

Energy availability (i.e., PET) and climate stability have a primary importance in determining the current spatial patterns of tetrapod μ-Diversity (Fig. 2). These two factors show a lower contribution when considering single taxonomic groups independently (Fig. S4-S7). Previous studies have shown that water–energy dynamics are important in describing species richness patterns^12, 34^. Our findings expand these previous findings to phylogenetic and functional aspects of diversity, providing a first comprehensive overview of a multifaceted diversity index and its components. Higher μ-Diversity in areas with high energy availability can be explained by higher resource availability, which in turn has promoted greater species packing (i.e., more species coexist with narrower niches) and larger population sizes, eventually lessening extinction rates^35^. Amphibians are the only group displaying a unimodal relationship with PET (Fig. S7), which is expected because of their high dependence on water^36^ and also because PET tends to increase towards drier environments, irrespective of their water-balance^37^. Furthermore, our results indicate that climate stability promotes higher diversity, probably through the combination of lower extinction rates and higher levels of speciation^38^ occurring also at a larger spatial scale. There is compelling evidence of higher extinction rates in climatically unstable regions, especially for those taxonomic groups with poorer dispersal abilities (e.g., amphibians)^39, 40^.

**Fig. 2.**
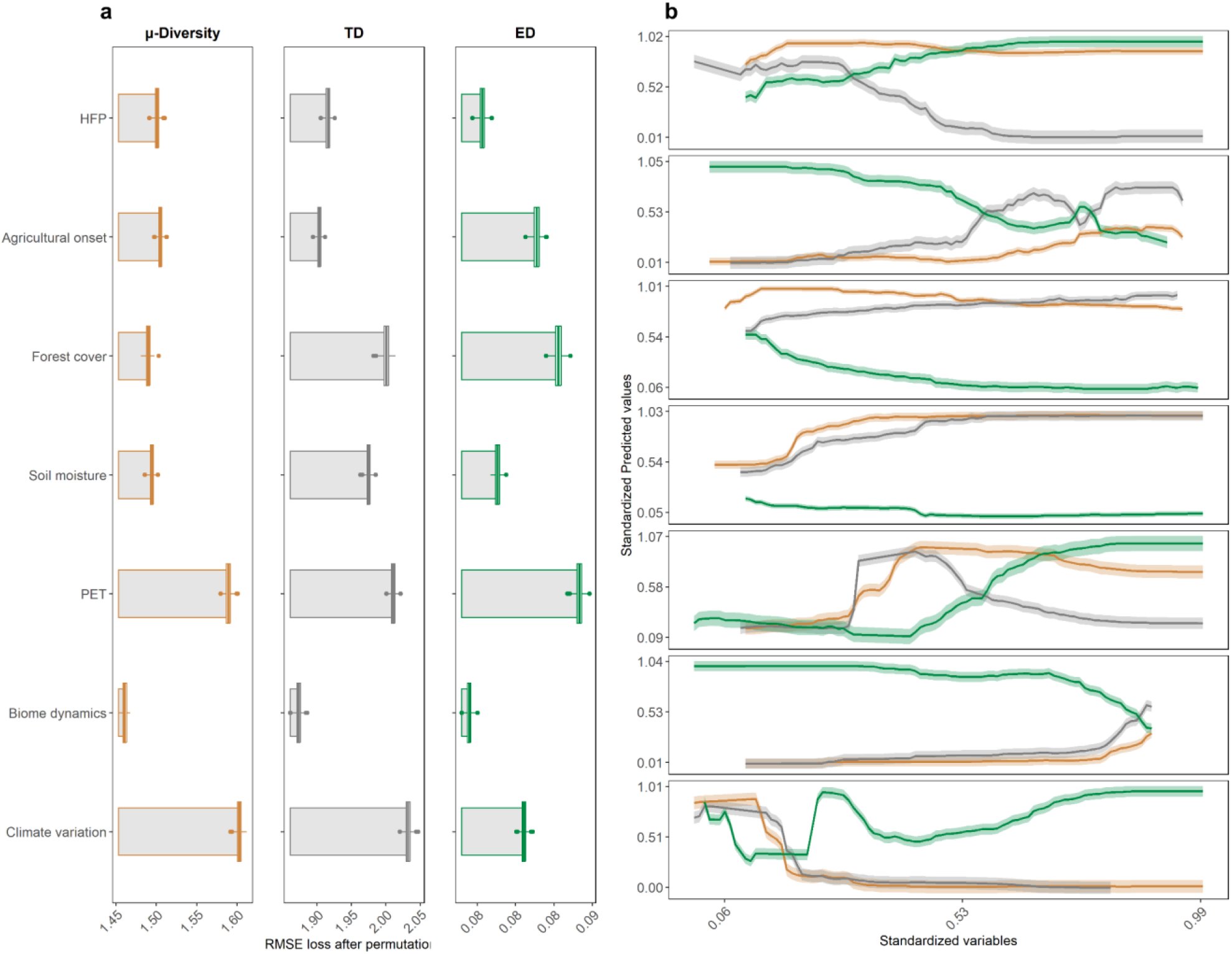
**a**, Variable importance obtained through Root Mean Square Error (RMSE) loss after permutations and **b**, marginal effects (mean ± 2 SE) of the different predictors of the random forest model using tetrapod μ-Diversity as response variable. Boxplots represent the values of RMSE loss in each permutation (*N* = 500) considering μ-Diversity, taxonomic diversity (TD) and ecological distinctness (ED). Climate stability represented the average rate of change of climate since Last Glacial Maximum (expressed in °C/century). Biome dynamic described the variation in biome patterns over the last 140 ka expressed using the Shannon index. PET, Soil moisture and Forest cover represented Potential Evapotranspiration, moisture in the soil and global forest cover updated to 2019, respectively. Agricultural onset expresses the year of onset of intensive agriculture *sensu* ref^20^, while *HFP* is the 2009 Human Footprint index. Orange colour denotes μ-Diversity, grey TD while green was used for ED.

Recent findings propose an early human role in triggering land-use changes and controlling vegetation change^21, 22, 41^. We discovered a pervasive role of past humans’ activities on current diversity patterns, especially for birds, mammals, and reptiles. For these groups, past human activities appear as important as or even more than present human impacts in predicting the broad-scale patterns (Fig. S4-S6); however, the direction of the relationship is variable among taxa and facets under consideration. Our results show that regions with an earlier onset of agriculture (e.g., Arabic peninsula, Turkey) tend to harbor lower tetrapod μ-Diversity (Fig. 2), suggesting that humans imprints might have influenced tetrapod diversity for millennia through legacy effects of past land use modifications^20, 21^. Recent findings suggest an acceleration in vegetation change during the Late Holocene (∼4.6 to 2.9 ka)^21^, which is in agreement with the onset of the Neolithic revolution (i.e., the shift from a society characterized by hunter-gatherers to one composed by farmers)^42^. Consequently, the expansion of pastoralism and agriculture might have triggered a gradual conversion of natural ecosystems via forest clearances and widespread use of fire (but see^43^). Moreover, domesticated animals might have started competitive interactions with local fauna, which, along with opportunistic hunting by humans, could have contributed to deplete local biodiversity^44^ across all diversity facets.

### Future global change threatens μ-Diversity in tropical and subtropical areas globally

We assessed how current tetrapod diversity might be threatened by future global change by estimating the magnitude of future climate and land use changes across the global distribution of μ-Diversity. Our results show that the areas hosting the highest μ-Diversity will be disproportionally exposed to global change by 2050. Exposure to global change is particularly high in tropical and subtropical regions of tropical Africa, South America, Indian subcontinent and South East Asia as well as Eastern Australia (Fig. 3, Figs. S8-S12 show exposure including the direction of change while Figs. S13-S18 in absolute terms, Table S4). Regardless of the scenario, about 70% of the areas hosting high tetrapod μ-Diversity such as Central and Western Africa or Borneo in South East Asia are expected to suffer dramatic changes in precipitation regime, land use and vegetation (see 3^rd^ and 4^th^ quartile in Fig. S14). By contrast, changes in temperature will primarily affect areas hosting lower tetrapod μ-Diversity that are mostly located in North America and Eurasia (1^st^ and 2^nd^ quartile). Especially in these regions where μ-Diversity is lower, an annual temperature change above 2 °C might translate in potential abrupt changes in local assemblages by 2050^2^ due to a more severe exposure to extreme thermal events^3^. Moreover, assemblages in these regions might be endowed of lower ecological redundancy with respect to the ones displaying higher μ-Diversity^45^, which makes them potentially less resistant to warmer climatic conditions^3^. For instance, among ectotherms^46^ (Fig. S17-S18), about 50% of the areas with lower μ-Diversity are going to be vastly exposed to rising temperature and precipitation change in the less sustainable scenario (e.g., increased droughts in the Amazonia and in the Mediterranean Basin), potentially causing overheating and dehydration, eventually hampering potential migrations towards more suitable climate^47^.

**Fig. 3.**
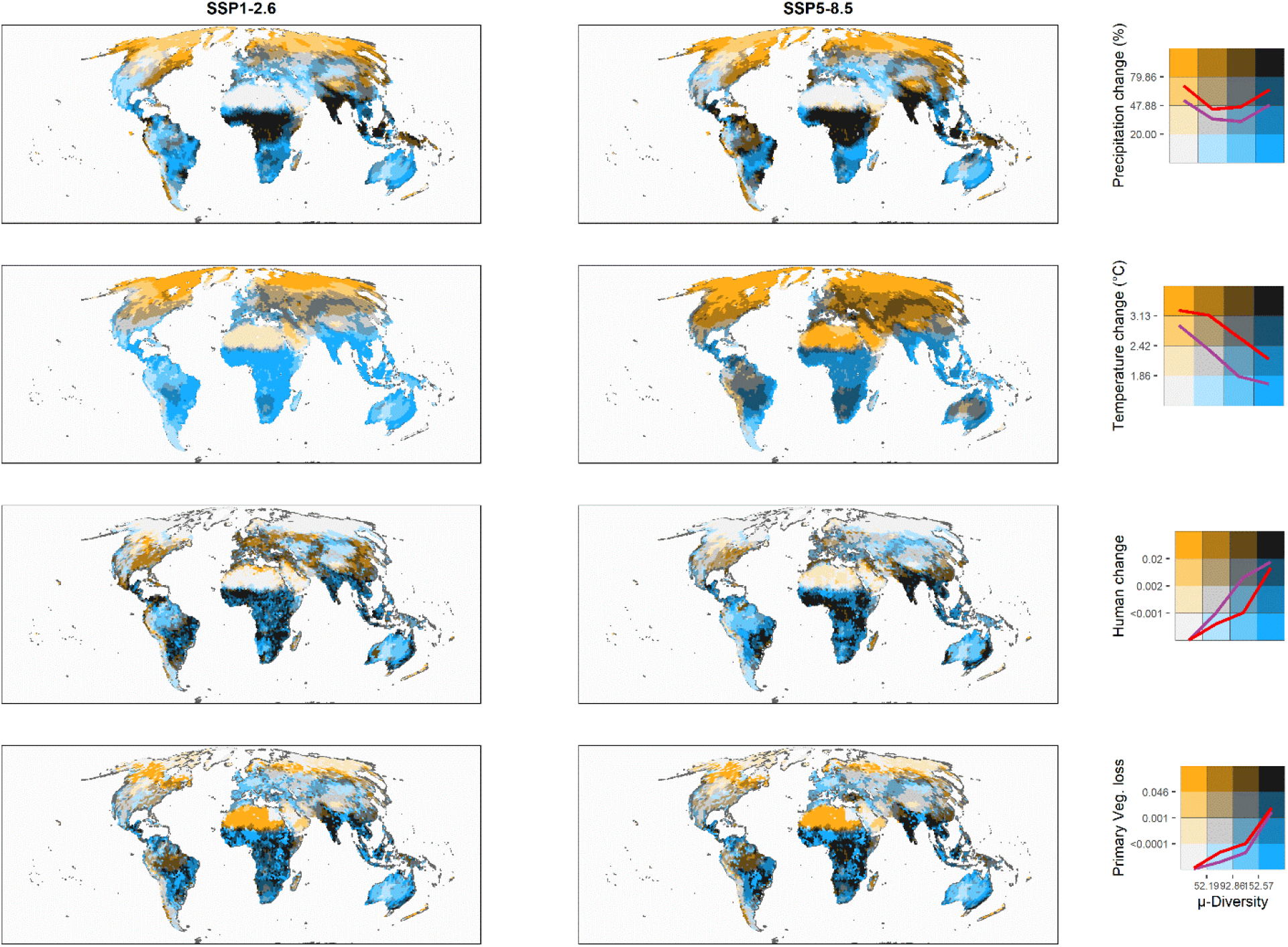
The areas hosting the highest levels of tetrapod μ-Diversity will be disproportionally exposed to projected global change. Bivariate maps showing the spatial matches between tetrapod μ-Diversity, going from light (low μ-Diversity) to dark blue (high μ-Diversity) along the x-axis of the legend, and the predicted absolute change (i.e. exposure) for each climate and land-use variable (from light to dark yellow along the y-axis). All variables are expressed in quartiles, with darker colors indicating areas of high μ-Diversity that will be highly exposed to the considered component of global change. Here we report only the two most extreme Tier 1 scenarios, spanning from a sustainability scenario based on green economy (i.e., “Taking the Green Road”, SSP1-2.6) to a high emission scenario mainly based on fossil fuel exploitation (i.e., “Fossil-fueled Development”, SSP5-8.5; see Fig. S13 for the intermediate scenarios). Colored lines in the legend connect the median value of the corresponding component of global change for each quartile of μ-Diversity, both for SSP1-2.6 (purple line) and SSP5-8.5 (red line), revealing how, irrespective of the scenario under consideration, cells with higher μ-Diversity are mostly exposed to precipitation change, human change and primary vegetation loss, while temperature change is affecting areas with lower μ-Diversity.

The synergistic interactions between anthropogenic climate and land-use change might exacerbate their effects on global biodiversity. The overarching importance of land-use change alone will probably increase the number of species under imminent imperilments^48^. The degradation of natural habitats caused by human encroachment strongly affects wildlife^4^, causing contractions in species ranges^49^ and even modifications in their behaviors^50^. Exposure of areas of high μ-Diversity to increased human pressure are particularly important in India and Eastern Africa, where the expected general increase in human population will probably increase the pressure on vulnerable tetrapod species^33^.

We show that combining different facets of diversity brings a more exhaustive evaluation of the overall diversity of life, which is essential to better frame conservation efforts in a world under accelerated environmental changes. Even though we found an overarching importance of energy availability and climate stability for tetrapod diversity, we detected a strong imprint of humans dating back at least to mid-Holocene, especially in mammals and amphibians, which seems also to be extremely sensitive to current human impacts. These anthropogenic impacts are likely to increase in the near future, particularly in the parts of the world that host the highest diversity of tetrapods, which are also disproportionally exposed to mid-21^st^ global change.

## Methods

### Species spatial distribution and environmental data

We obtained expert-verified range maps of 23,848 tetrapod species from the International Union for Conservation of Nature (IUCN)^51^. Even though these maps might underestimate the complete extent of occurrence of the species, especially in poorly surveyed regions^52^, they currently represent the best available source. After excluding marine mammals, range maps were intersected with a hexagonal equal-area grid cells distributed at regular distances across the global mainland (cell area of 23,322 km^2^) using the ‘dggridR’^53^ R package. We chose this resolution because it is close to the finest resolution justifiable for using global data without incurring in false presences^54^. Realm’s limits were downloaded from^55^ and overlaid with grid cells using the R package ‘sp’^56^. Species names were standardized using the Global Biodiversity Information Facility (GBIF) Backbone Taxonomy^57^ with the R package ‘taxize’^58^.

For each grid cell, we computed predictors depicting spatiotemporal effects of past climate change/biogeography history and current ecosystem features along with past and current anthropogenic variables. Climate stability since Last Glacial Maximum (ca. 20 ka) was estimated as the median rate of change during the time-series expressed in °C/century^59^, so that larger values indicate less stability and vice versa. Biome dynamic described the dynamic in biome patterns over the last 140 ka. In more details, biome extents and locations were gathered from^60^, which was inferred through a process-based model (LPJ-GUESS) simulating the global vegetation starting from 140 ka BP. Biome dynamic was obtained computing the Shannon index for each grid cell across the time series. As environmental features, we selected three variables encompassing factors which have been shown to be important drivers especially for tetrapod diversity, namely Potential Evapotranspiration (PET) as a proxy of ambient energy^61^, forest cover of habitat characteristics and soil moisture as a proxy of water availability^62, 63^. Gridded data of PET and soil moisture were obtained from TerraClimate^64^, while forest cover updated to 2019 was retrieved from Copernicus Global Land Cover products^65^. Past human influence was approximated as the year of onset of intensive agriculture (i.e., all types of cultivation characterized by a continuity in time) sourced from the ArchaeoGLOBE Project^20^. To depict the spatial distribution of the current human pressure across the globe, we used the 2009 Human Footprint index (HFP2009)^33^, which reports for each grid cell a measure of the intensity of eight metrics of human pressure (i.e., human population density, roads, railways, navigable waterways, built environments, crop land, pasture land, night-time lights), weighted based on the relative human pressure on that cell. Multicollinearity among predictors was checked ensuring that Spearman’s correlation coefficient (ρ) was < |0.7|^66^ (Table S5).

### Functional traits

Functional trait data for the different groups were collected using public databases from different sources, using the same trait data available in Carmona et al.^45^. The selected traits reflect the ecological strategies of species in relation to reproduction, longevity and size (i.e., life-history traits; Table S6).

#### Mammals, reptiles and birds

Data were retrieved from Amniote database^67^, which include traits for 4,953 species of mammals, 6,567 species of reptiles, and 9,802 species of birds. Specifically, this database contains information of 29 life history traits, of which we selected a subset of traits with information available for at least 1000 species. For mammals, eight traits were chosen: longevity (*long*, years), number of litters per year (*ly*), adult body mass (*bm*, g), litter size (*ls*, number of offspring), weaning length (*wea*, days), gestation length (*gest*, days), time to reach female maturity (*fmat*, days), and snout–vent length (*svl*, cm). For birds, we selected the following traits: number of clutches per year, adult body mass (*bm*, g), incubation time (*gest*, days), clutch size (*ls*, number of eggs), longevity (*long*, years), egg mass (*em*, g), snout-vent length (*svl*, cm), and fledging age (*fa*, days). Regarding reptiles, six traits were selected: number of clutches per year, longevity (*long*, years), adult body mass (*bm*, g), clutch size (*ls*, number of eggs), incubation time (*inc*, days), and snout-vent length (*svl*, cm).

#### Amphibians

Functional trait data of 6,776 species of amphibians were retrieved from AmphiBIO database ^68^. We selected four traits that mirror similar information as the one collected for the other three groups (i.e. traits related to body size, pace of life and reproductive strategies): age at maturity (*am*, years), body size (*bs*; measured in Anura as snout-vent length – *SVL* – and in Gymnophiona and Caudata as total length in mm); maximum litter size (*ls*, number of individuals); and offspring size (*os*, mm).

### Phylogenies

Phylogenies for each group were downloaded from published papers^69–72^. Species absent from the phylogeny were manually added to the root of their genus using the R package ‘phytools’^73^. Since for mammals and birds multiple phylogenetic trees were available, we computed a maximum clade credibility tree (MCC) for these groups using the ‘phangorn’^74^ R package. To assess the reliability of the information contained in the MCC, we performed a simulation where we correlated cell-based phylogenetic diversity (PD^5^, i.e., the sum of branch length between the root node and tips for the subtree comprising all species in the assemblage). PD obtained from this MCC with those obtained with 100 phylogenies randomly selected from the original posterior distribution. This test proved that using the MCC tree is unlikely to affect the computation of PD (Fig. S19).

### Construction of the functional spaces, trait imputation, and sensitivity analysis

Due to the presence of gaps in the functional trait data (see^45^ for more details about the completeness of trait data), we first imputed missing traits for each group using ‘missForest’^75^ R package (step 1). We used a Random Forest algorithm to impute trait data taking advantage also of the phylogenetic relationships among species (included as the first 10 components of a PCoA based on the phylogenetic dissimilarity among pairs of species), as described in Penone et al.^76^. We computed two Principal Component Analysis (PCA) (one with and one without phylogeny), determined their dimensionality (two dimensions in all cases) and retained the scores of the species in each of these axes as indicators of their position in the trait space (step 2). To further validate the imputation process, we performed a sensitivity analysis similar to the one performed in Carmona et al.^45^ (step 3), but repeating the imputation process with and without using phylogenetic information to evaluate the effect of considering phylogeny in the imputation process. Specifically, we simulated missing traits (100 repetitions) starting from a subset of species with complete trait data. We then randomly selected 10% of species assigning them the structure of missing values of the same number of randomly selected species from the subset of species with missing trait values. Once we obtained this simulated dataset (with artificial missing data, plus all original missing data, so that the procedure is very conservative), we projected the positions of the species in the PCA space performed in step 2. We then compared the position of the species to which we added artificial missing values after this “projection” with their “real” position. For only the species with artificial missing values, we evaluated the normalized root mean square error (NRMSE) between the original position in the functional space and the position calculated after trait imputation, expressed as the relative range of trait values in the corresponding PC axis. Our sensitivity analysis showed that the imputation process is reliable in retrieving the positions of species in the functional space showing i) high accuracy of the imputations with an average NRMSE of 7 ± 5% (mean ± SD) across taxonomic groups and PC axes, and ii) the procedure that includes phylogeny halves the NRMSE on average with respect to the imputation realized with traits information only (Fig. S20). Furthermore, considering that phylogenetic information was used only to improve the precision in placing the species onto the functional space, and accounting for that two species can be closely related under a functional point of view while being distant from an evolutionary perspective, this makes us relatively confident about the risk to artificially influence the overlap between functional and phylogenetic metrics. For all these reasons, we used the PCA obtained using the imputation with phylogeny information in the rest of the analysis. Once we imputed traits, we removed from the database extinct species and species totally lacking evolutionary, trait or spatial data, thus leaving 17,341 species for the calculation of the diversity metrics (*N* = 3,912 for mammals, *N* = 3,239 for amphibians, *N* = 3,338 for reptiles and *N* = 6,852 for birds; see Table S1 for more details) with an average taxonomic completeness of 0.83 (range 0.42-1.00).

### Calculation of diversity metrics

For each grid cell of mammals, birds, amphibians and reptiles, we calculated TD measured as species richness, PD using Faith’s PD^5^, and FD which was expressed as functional richness (i.e., the amount of the functional space occupied by an assemblage ^77^). PD was computed using the ‘caper’^78^ R package. To estimate FD, we first calculated the functional space as described above, then by means of TPD framework^77^ and ‘TPD’ and ‘ks’ R packages^79, 80^, we estimated cell-based functional richness. Since both PD and FD are strongly dependent on species richness, we performed null model simulations to break this relationship^81^ and compute standardized effect sizes (SES) as SES = (Metric_obs_-mean(Metric_null_))/SD(Metric_null_). To obtain the null distribution, we randomized 1000 times the community composition of each cell preserving marginal totals using the quasiswap algorithm in the R package ‘vegan’^82^. Positive SES values indicate that the considered variable (i.e., FD or PD) is higher than expected considering the number of species present in a grid cell, and vice versa. We allowed the algorithm to shuffle species across the whole world avoiding geographical stratification since we were more interested in obtaining worldwide comparisons which reflect the overall global diversity in our dataset. In addition, in order to mitigate the effect of random variation, we considered only the grid-cells hosting more than 3 species for further analysis. We thus averaged sesPD and sesFD to calculate the Ecological Distinctiveness – ED – a metric which reflects the amount of functional and phylogenetic information contained in a given grid cell. Specifically, after having computed sesPD and sesFD, we centered and scaled to unit variance sesPD and sesFD in order to obtain comparable ranges of variation across taxonomic groups, and then we re-scaled these values between 0−1 by comparing them with a cumulative normal distribution function with mean = 0 and standard deviation = 1 to remove the effect of extreme values. Finally, the latter were averaged to compute ED for each grid cell. μ–Diversity for each taxon was calculated independently in each grid cell by multiplying TD for its related ED. The calculation of μ-Diversity is conceptually analogous to an effective diversity whereby

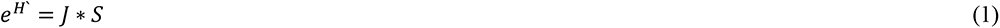

where H’ is Shannon-Wiener index, *J* is Hill’s evenness ratio and S is the total number of species in a given assemblage. In our case, S still represents the total number of species in a given assemblage whilst the evenness is expressed in terms of ecological distinctiveness. In this way, two assemblages having the same species richness but different values of ED (e.g., ≈ 1 or ≈ 0) will have a μ-Diversity that will be closer to the species richness in case of ED ≈ 1, otherwise observed species richness will be penalized proportionally using ED as a weight. Finally, tetrapod μ-Diversity was obtained as the overall sum of the μ-Diversity of each taxonomic group. To test for spatial associations among original diversity facets (i.e., SR, sesPD, sesFD), we computed the Pearson’s correlation (*r*) by using a modified t-test of spatial association^83^ implemented in the ‘SpatialPack’ R package^84^.

### Drivers of diversity

We used Random Forest (RF) to assess the relationship between μ-Diversity, TD and ED and their hypothesised drivers (i.e., past climate change/biogeography history, current ecosystem features along with past and current anthropogenic variables). We build five random forest models (one for tetrapod plus one for each individual taxonomic group) studying the response of μ-Diversity to current and past environmental and human predictors using the framework provided in the ‘ml3’^85^ and ‘mlr3spatiotempcv’^86^ R packages. We started building trees using the following hyperparameters: number of trees (*ntree*) = 500, number of variables randomly sampled at each split (*mtry*) = 1, minimum size of terminal nodes (*min.node.size*) = 1, and sample fraction (*sample.fraction*) = 0.6, which were later tuned using the ‘paradox’^87^ R package. Variable importance was determined by measuring the mean change in a loss function (i.e., Root Mean Square Error – RMSE) after variable permutations (*N* = 500) using ‘DALEX’ R package^88^. This method assumes that if a variable is relevant for a given model, we expect a worsening in model’s performance after randomly permuting its values (see^89^ for more technical details). In other words, this method assesses variable importance as the loss in explanatory ability of the model when that variable is randomized. We also displayed marginal effects of different predictors by using partial dependence plots computed with the ‘iml’^90^ R package. We computed three models for each taxonomic group using μ-Diversity, TD and ED as response variables. For tetrapods, μ-Diversity and TD were obtained by summing the values of each taxon, whereas ED was calculated as the arithmetic mean of each taxon ED to ensure that its value remains between 0 and 1.

### Spatial cross validation

Failing to account for spatial autocorrelation processes in ecology might lead to biased conclusions^91, 92^ or to an overoptimistic evaluation of model predictive power^93, 94^. For this reason, we performed an internal spatial cross-validation (spCV). We created five spatially disjointed subsets (i.e., partitions) which try to maximize the spatial distance between the training and validation sets so that these sets were more distant than they would have been using random partitioning^95^. To perform the spCV, we used a nested resampling approach as described in ref.^96^, where outer resampling evaluated model performance while inner resampling performed tuning of model hyperparameters for each outer training set. Because nested resampling is computationally expensive, we selected 5-folds with 5 repetitions each to reduce the variance introduced by partitioning in outer resampling and 5-folds in inner resampling coupled with 50 evaluations of model settings.

### Exposure to mid-21st global change

The future climate and land-use projections derive from the sixth phase of the Coupled Model Intercomparison Project (CMIP6). Climate data were sourced from Copernicus Climate Data Store (https://cds.climate.copernicus.eu/cdsapp#!/dataset/projections-cmip6?tab=overview), while land use data were sourced from the land use harmonization 2 (LUH2) dataset (51; https://luh.umd.edu/data.shtml). These harmonized climate and land-use projections combine alternative Shared Socioeconomic Pathways (SSP) and Representative Concentration Pathways (RCP) spanning from milder scenarios of sustainable growth and low global warming (SSP1-RCP2.6) to more extreme forecasts (SSP5-RCP8.5) of development based mainly on fossil-fuel exploitation^98^. Following the guidelines provided by the sixth assessment report of IPCC (https://www.ipcc.ch/report/ar6/wg1/#TS), climate change was expressed in terms of anomalies between a baseline period (1961–1990) and mid-21st century (2040–2069). We used a single model variant from a total of 24 General Circulation Models (GCMs) and for each GCM and scenario we calculated the difference (i.e. anomaly) between future climate (i.e. average of years 2040–2069) and the baseline period (i.e. average of years 1961–1990) obtained by each GCM’s historical run. Individual model anomalies were then resampled to 1.4° × 1.4° spatial resolution and multi-model ensemble means were computed for each future scenario. Land use change was computed as the difference in the frequency of land use categories between current conditions (year 2015) and 2050 under each future scenario. We considered in total 4 Tier 1 scenarios, namely SSP1-RCP2.6, SSP2-4.5, SSP3-7.0 and SSP5-8.5, and four variables: near-surface annual temperature change (°C), precipitation change (mm), change caused by human activities (e.g., human settlements, pastures, croplands) and primary vegetation loss (e.g., pristine habitats). For further details about data processing see ^99^. We used bivariate maps intersecting μ-Diversity with the maps of predicted anomalies in annual temperature and precipitation (1.4° × 1.4°) and land-use change (0.25° × 0.25°) at grid-cell scale considering both absolute and relative changes. To ease data visualization, the values of the variables in each grid-cell were previously classified in quartiles (i.e. 0–25, 25–50, 50–75, 75–100%). We further computed the Pearson’s correlation (r) between μ-Diversity and the predicted absolute changes described above by using a modified t-test of spatial association^83^ implemented in the ‘SpatialPack’ R package^84^.

## ACKNOWLEDGMENTS

We are grateful to Stefano Mammola for his comments provided on a previous version of the manuscript. Part of the analyses were carried out using the facilities of the High-Performance Computing Center of the University of Tartu. E.T., C.P.C., A.T., and M.P. were supported by the Estonian Research Council grants (MOBJD1030, PSG293, MOBERC40, PRG609, and PSG505) and the European Regional Development Fund (Centre of Excellence EcolChange).

## Author contributions

E.T. and C.P.C. co-led and designed the study. E.T., A.T. and C.P.C. extracted and prepared the data, S.T. provided the global change scenarios, E.T. performed the statistical analyses, M.P. and D.N.B. contributed to the interpretation of results. E.T. led the writing of the manuscript with inputs from all the co-authors who approved the submitted version.

## Data and materials availability

In case of paper acceptance, the data and R scripts needed to reproduce the figures in the main text will be made available in a public repository (e.g., figshare, zenodo). Additional data and scripts related to this paper will be available from the corresponding author upon request.

## Supplementary Information

### Figures and Tables

**Fig. S1.**
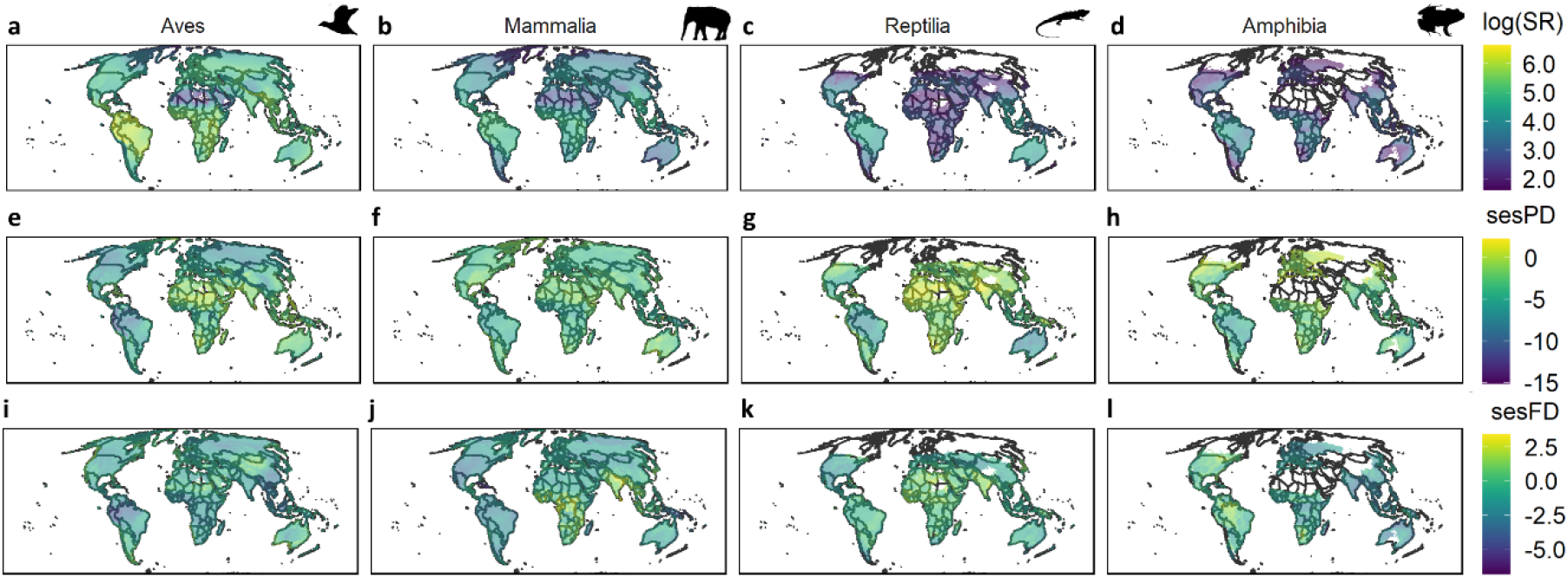
Global patterns of the taxonomic, phylogenetic and functional diversity of the four tetrapod groups. **a-e-i**) Aves, **b-f-j**) Mammalia, **c-g-k**) Reptilia, **d-h-l**) Amphibia. Taxonomic diversity was calculated as the number of species in each grid cell. sesPD is the standardized effect size of phylogenetic diversity (PD), while sesFD represents the standardized effect size of functional diversity (FD), expressed as functional richness. Standardized effect sizes (SES) were computed as follows: SES = (Metricobs−mean(Metricnull))/SD(Metricnull). Please note that species richness was *log (x+1)* transformed to improve data visualization. Silhouettes were retrieved from PhyloPic (www.phylopic.org).

**Fig. S2.**
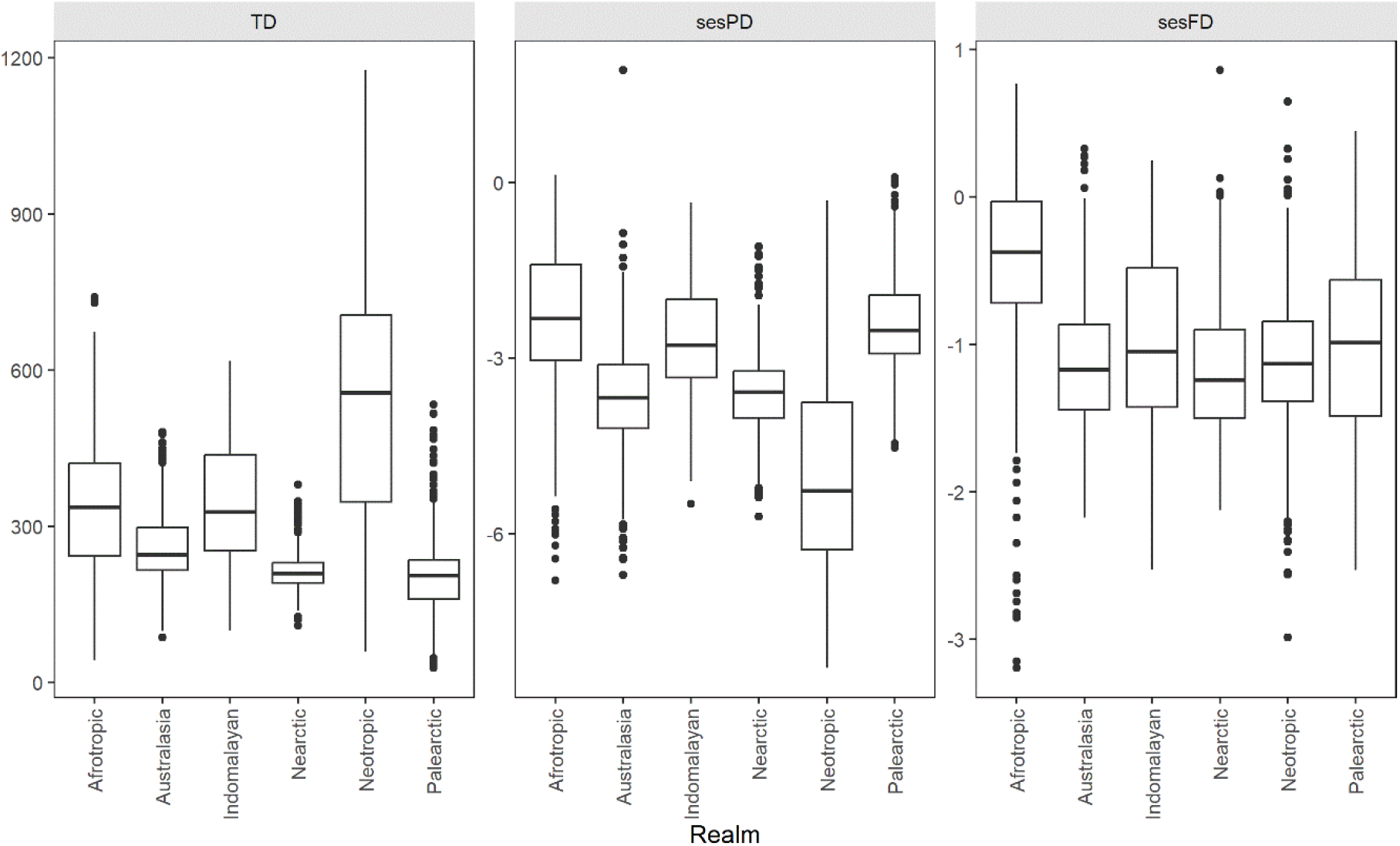
Boxplots showing the distributions of tetrapod species richness (TD), phylogenetic diversity (sesPD) and functional diversity (sesFD) for each realm. Please note that sesPD and sesFD represent standardized effect sizes (SES) of the original metric. SES were computed as follows: SES = (Metricobs−mean(Metricnull))/SD(Metricnull). SR was calculated as the sum of the number of species across taxonomic groups, while sesPD and sesFD are the mean values. Please note that diversity facets were computed only in the cells where all the three metrics were available (*N* = 4622).

**Fig. S3.**
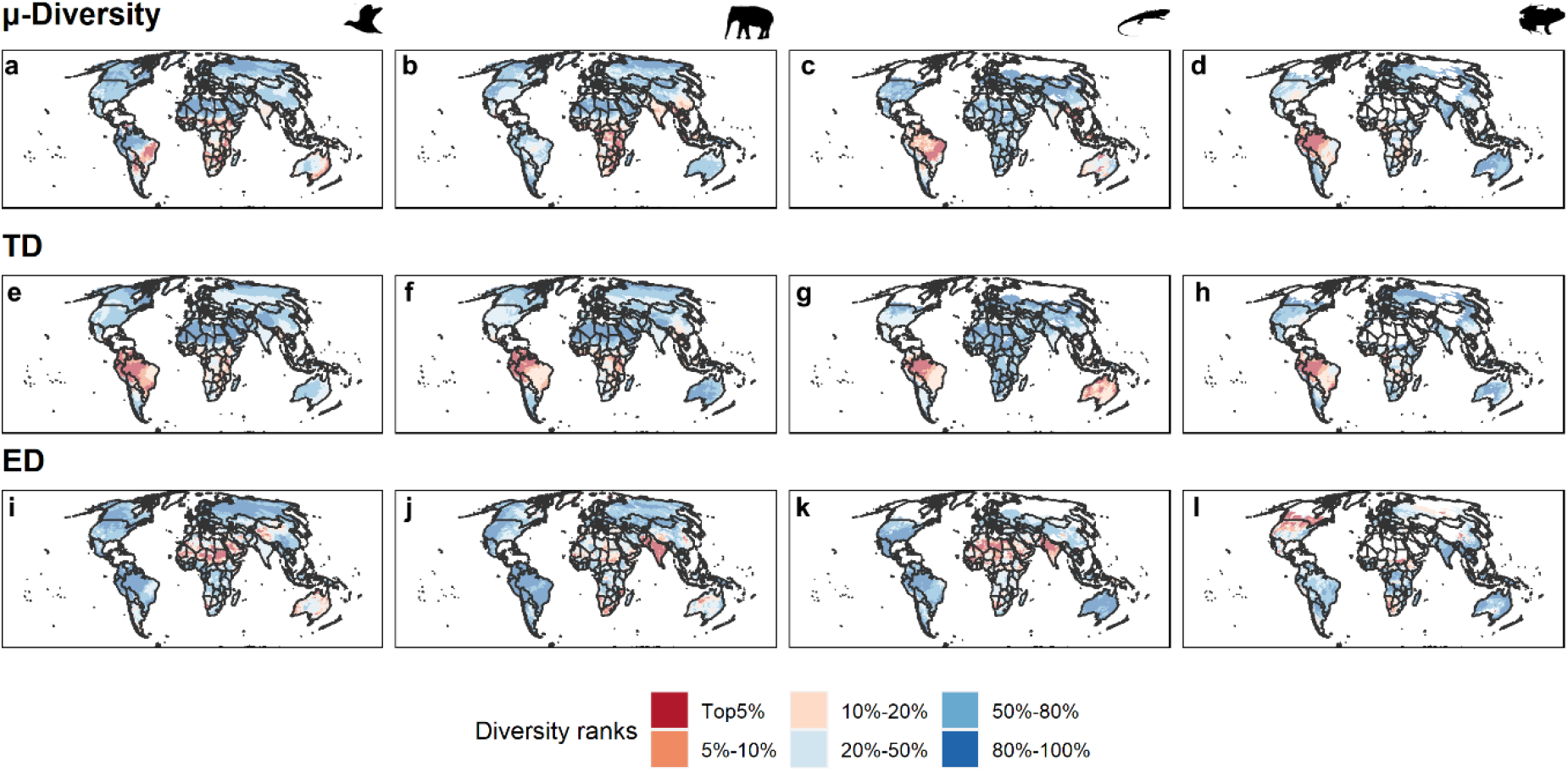
Global patterns of μ-Diversity, TD and ED. **a-e-i)** Aves, **b-f-j)** Mammalia, **c-g-k)** Reptilia, **d-h-l)** Amphibia. μ-Diversity was expressed weighting species richness by its ecological distinctiveness (ED) in each taxonomic group. TD is the species richness, while ED is the average between phylogenetic (sesPD) and functional diversity (sesFD) in each cell of each taxonomic group, after having scaled them between 0–1. Each diversity facet was ranked by the most (1–5) to least (80–100) diverse areas globally. The darkest brown tones denote the richest grid cells while blue tones indicate cold spots. Silhouettes were retrieved from PhyloPic (www.phylopic.org).

**Fig. S4.**
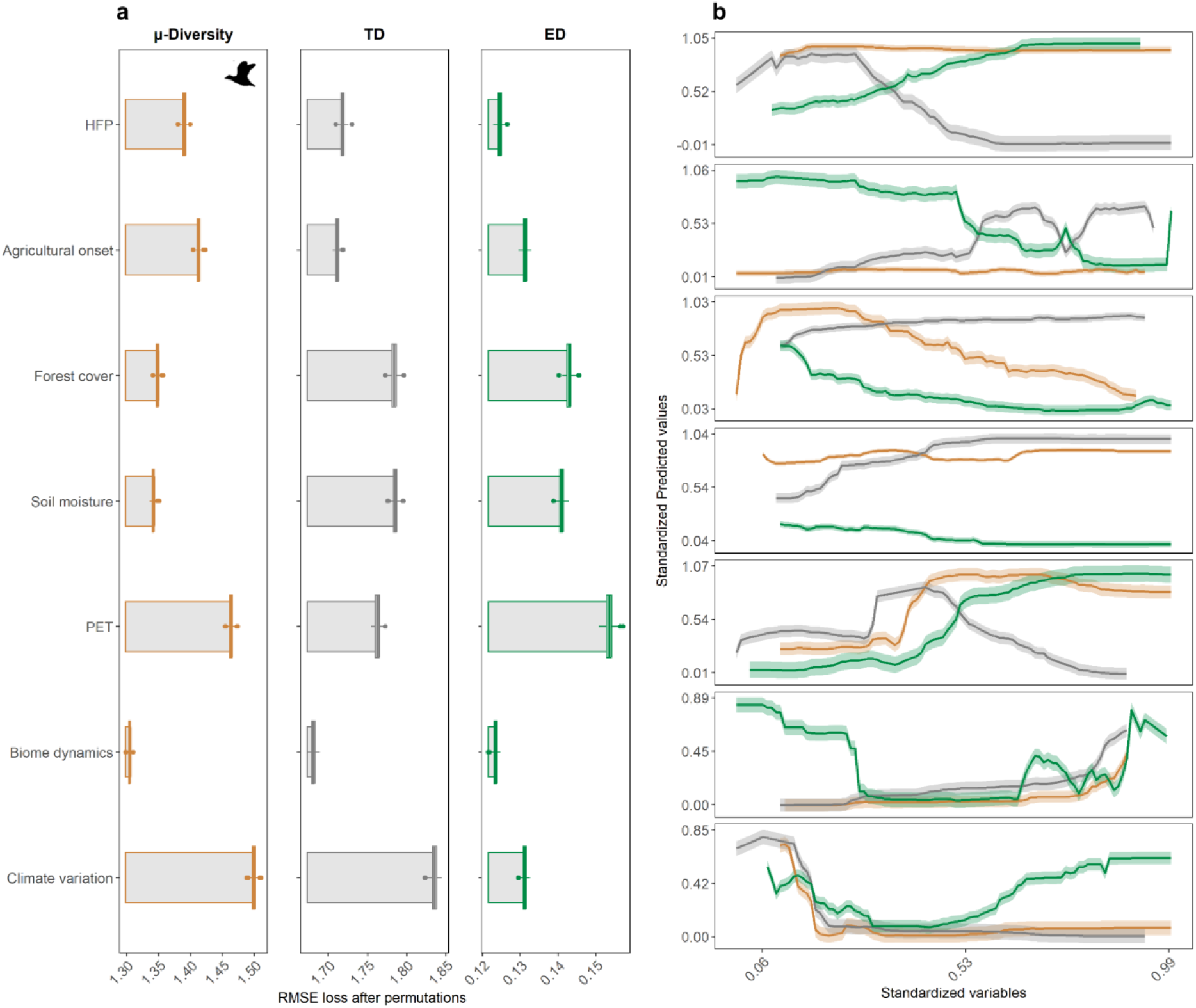
a) Variable importance ranked by the Root Mean Square Error (RMSE) loss after permutations and b) marginal effects (mean ± 2 SE) of the different predictors of the random forest model using avian μ-Diversity as response variable. Boxplots represent the values of RMSE loss in each permutation (*N* = 500) considering μ-Diversity, taxonomic diversity (TD) and ecological distinctness (ED). Climate stability represented the average rate of change of climate since Last Glacial Maximum (expressed in °C/century). Biome dynamic described the variation in biome patterns over the last 140 ka expressed using the Shannon index. PET, Soil moisture and Forest cover represented Potential Evapotranspiration, moisture in the soil and global forest cover updated to 2019, respectively. Agricultural onset expresses the year of onset of intensive agriculture *sensu* ref^20^, while *HFP* is the 2009 Human Footprint index. Orange color denotes μ-Diversity, grey TD while green was used for ED. Silhouettes were retrieved from PhyloPic (www.phylopic.org).

**Fig. S5.**
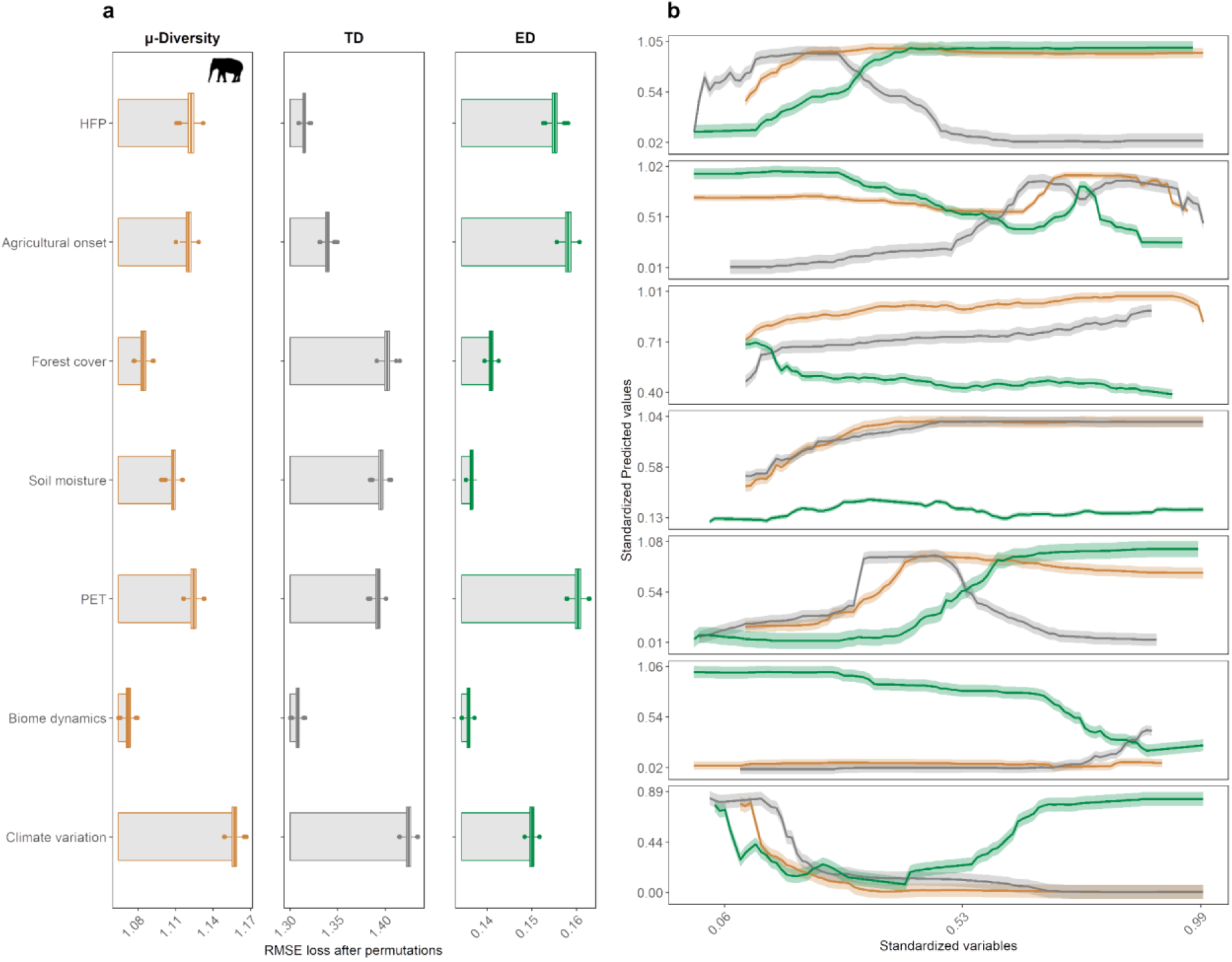
a) Variable importance ranked by the Root Mean Square Error (RMSE) loss after permutations and b) marginal effects (mean ± 2 SE) of the different predictors of the random forest model using mammal μ-Diversity as response variable. Boxplots represent the values of RMSE loss in each permutation (*N* = 500) considering μ-Diversity, taxonomic diversity (TD) and ecological distinctness (ED). Climate stability represented the average rate of change of climate since Last Glacial Maximum (expressed in °C/century). Biome dynamic described the variation in biome patterns over the last 140 ka expressed using the Shannon index. PET, Soil moisture and Forest cover represented Potential Evapotranspiration, moisture in the soil and global forest cover updated to 2019, respectively. Agricultural onset expresses the year of onset of intensive agriculture *sensu* ref^20^, while *HFP* is the 2009 Human Footprint index. Orange color denotes μ-Diversity, grey TD while green was used for ED. Silhouettes were retrieved from PhyloPic (www.phylopic.org).

**Fig. S6.**
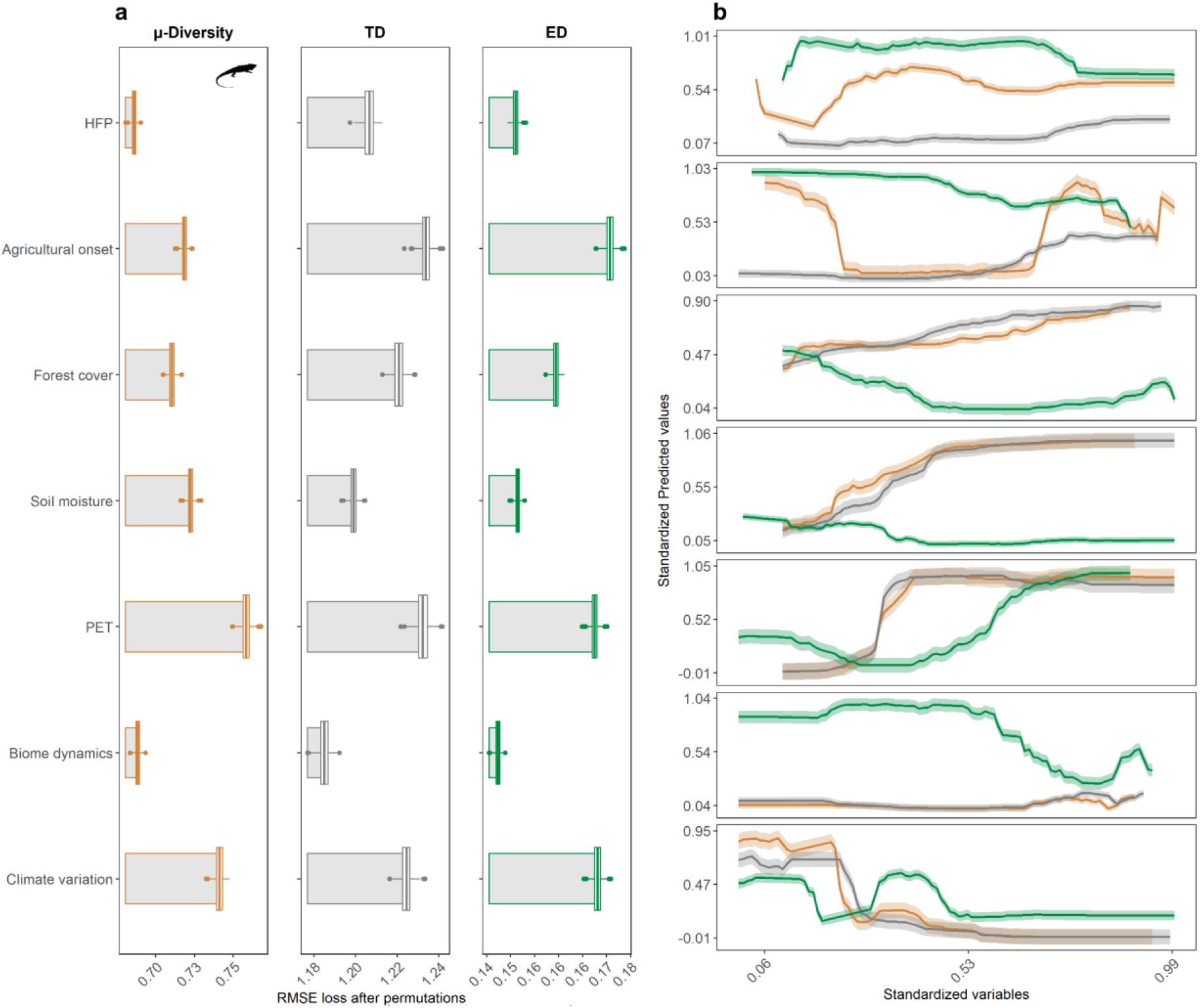
a) Variable importance ranked by the Root Mean Square Error (RMSE) loss after permutations and b) marginal effects (mean ± 2 SE) of the different predictors of the random forest model using reptile μ-Diversity as response variable. Boxplots represent the values of RMSE loss in each permutation (*N* = 500) considering μ-Diversity, taxonomic diversity (TD) and ecological distinctness (ED). Climate stability represented the average rate of change of climate since Last Glacial Maximum (expressed in °C/century). Biome dynamic described the variation in biome patterns over the last 140 ka expressed using the Shannon index. PET, Soil moisture and Forest cover represented Potential Evapotranspiration, moisture in the soil and global forest cover updated to 2019, respectively. Agricultural onset expresses the year of onset of intensive agriculture *sensu* ref^20^, while *HFP* is the 2009 Human Footprint index. Orange color denotes μ-Diversity, grey TD while green was used for ED. Silhouettes were retrieved from PhyloPic (www.phylopic.org).

**Fig. S7.**
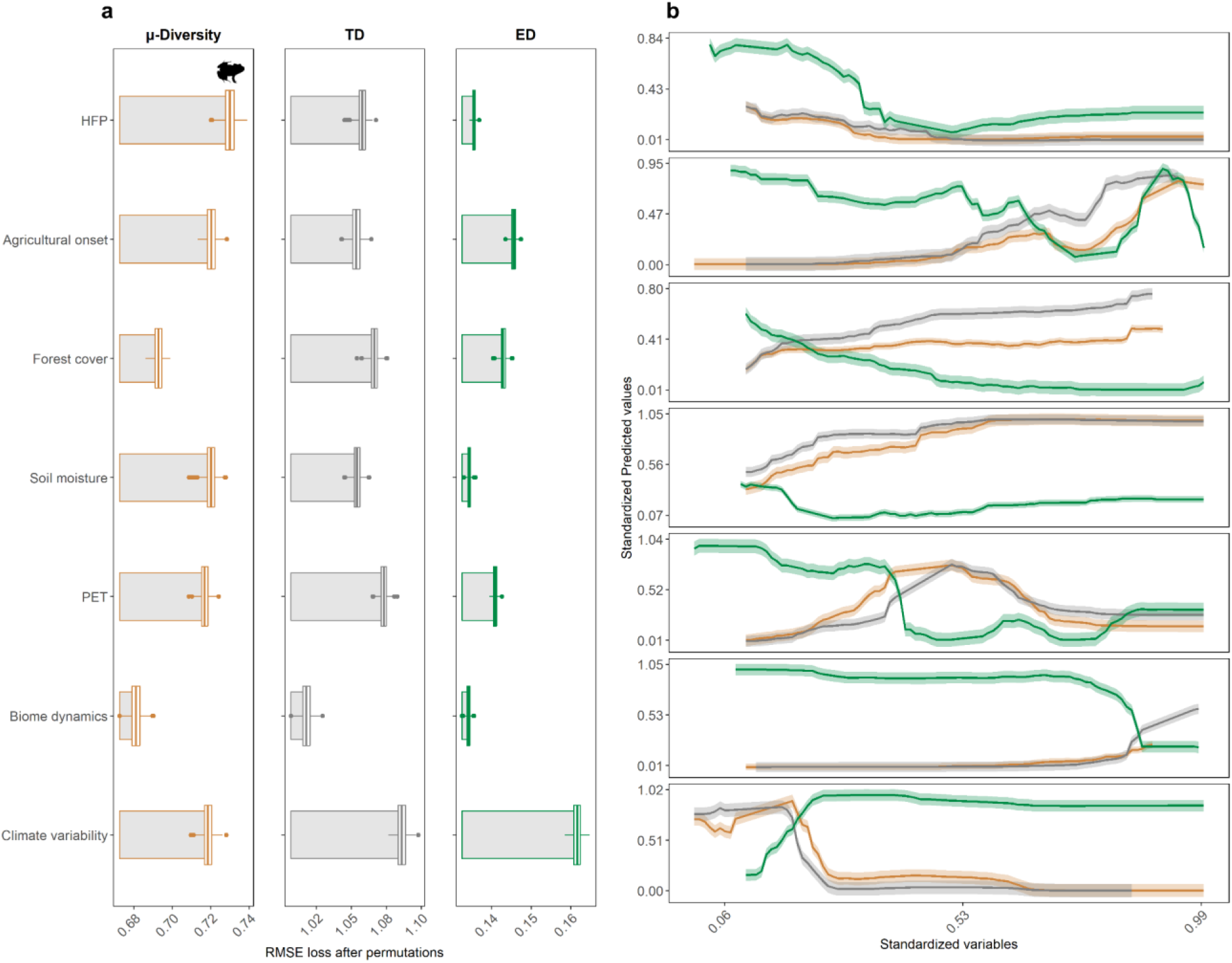
a) Variable importance ranked by the Root Mean Square Error (RMSE) loss after permutations and b) marginal effects (mean ± 2 SE) of the different predictors of the random forest model using amphibian μ-Diversity as response variable. Boxplots represent the values of RMSE loss in each permutation (*N* = 500) considering μ-Diversity, taxonomic diversity (TD) and ecological distinctness (ED). Climate stability represented the average rate of change of climate since Last Glacial Maximum (expressed in °C/century). Biome dynamic described the variation in biome patterns over the last 140 ka expressed using the Shannon index. PET, Soil moisture and Forest cover represented Potential Evapotranspiration, moisture in the soil and global forest cover updated to 2019, respectively. Agricultural onset expresses the year of onset of intensive agriculture *sensu* ref^20^, while *HFP* is the 2009 Human Footprint index. Orange color denotes μ-Diversity, grey TD while green was used for ED. Silhouettes were retrieved from PhyloPic (www.phylopic.org).

**Fig. S8.**
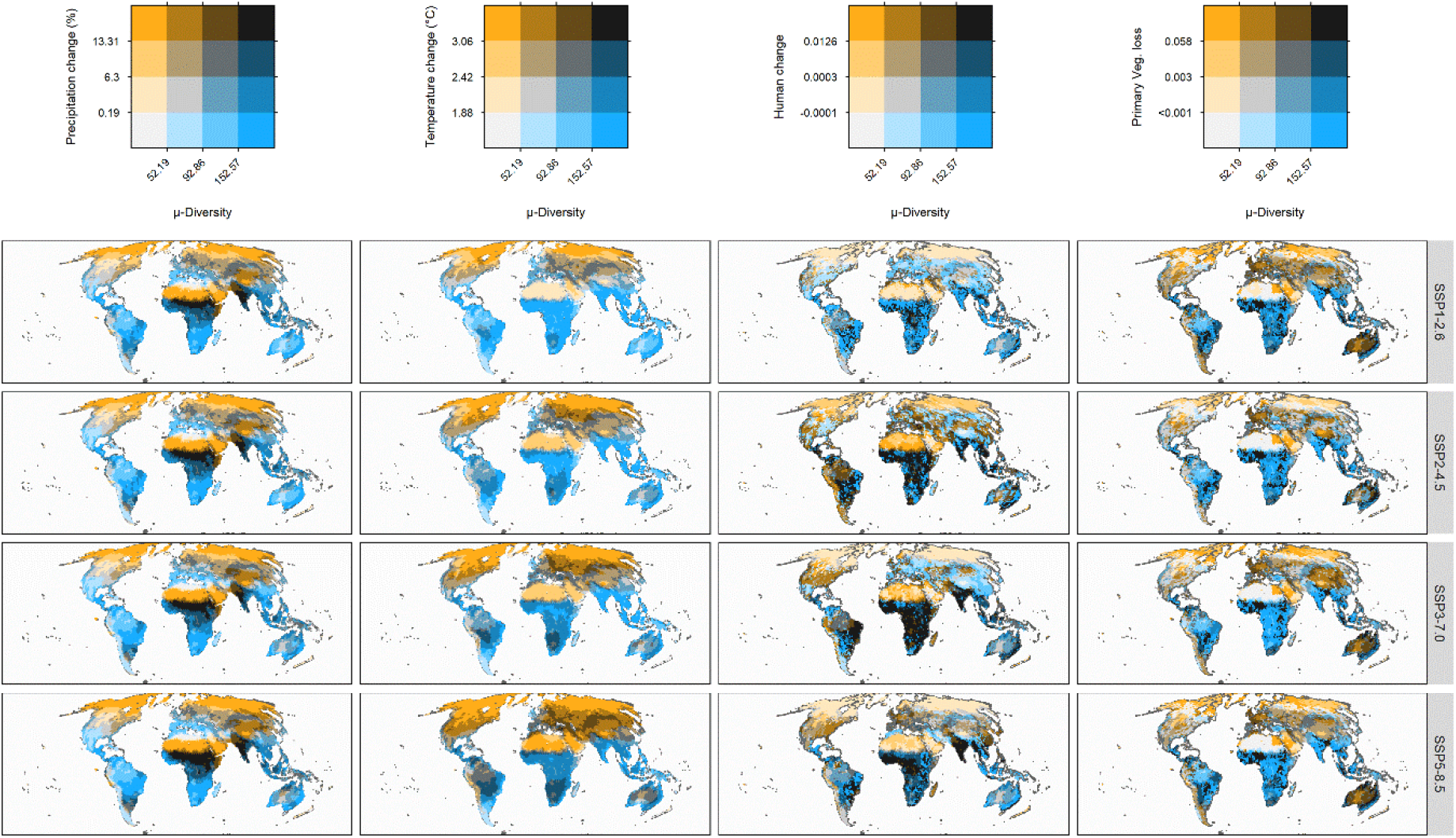
Exposure of tetrapod μ-Diversity to projected global change across four socioeconomic future scenarios. The y-axis indicates the predicted relative change (i.e. exposure) for each climate and land-use variable, while x-axis represents μ-Diversity. Both axes were expressed in quartiles, darker tones indicate larger exposure in relation to μ-Diversity. We considered four Tier 1 scenarios spanning from a sustainability scenario based on green economy (i.e., “Taking the Green Road”, SSP1-2.6) to a high emission scenario mainly based on fossil fuel exploitation (i.e., “Fossil-fueled Development”, SSP5-8.5).

**Fig. S9.**
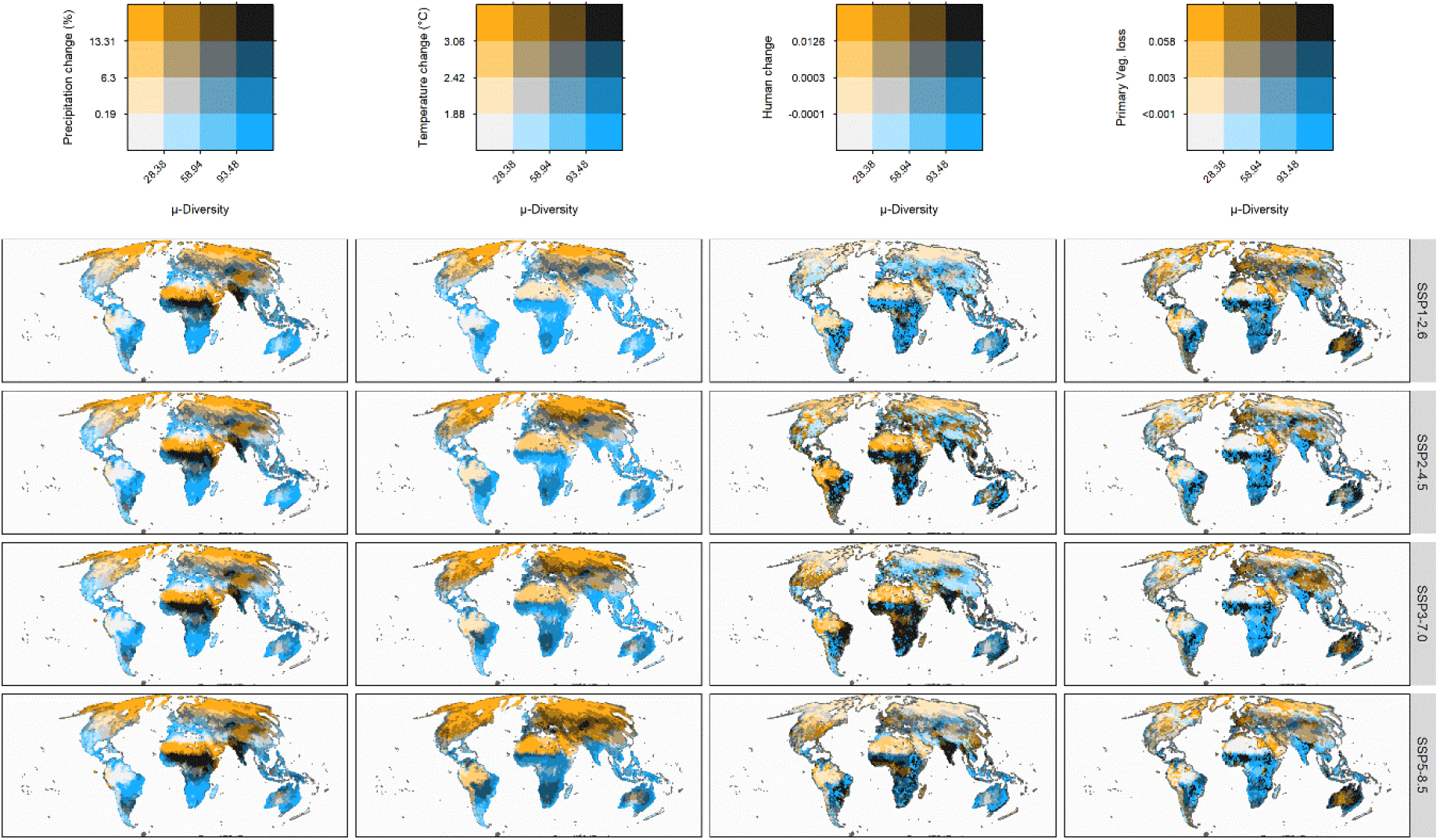
Exposure of avian μ-Diversity to projected global change across four socioeconomic future scenarios. The y-axis indicates the predicted relative change (i.e. exposure) for each climate and land-use variable, while x-axis represents μ-Diversity. Both axes were expressed in quartiles, darker tones indicate larger exposure in relation to μ-Diversity. We considered four Tier 1 scenarios spanning from a sustainability scenario based on green economy (i.e., “Taking the Green Road”, SSP1-2.6) to a high emission scenario mainly based on fossil fuel exploitation (i.e., “Fossil-fueled Development”, SSP5-8.5).

**Fig. S10.**
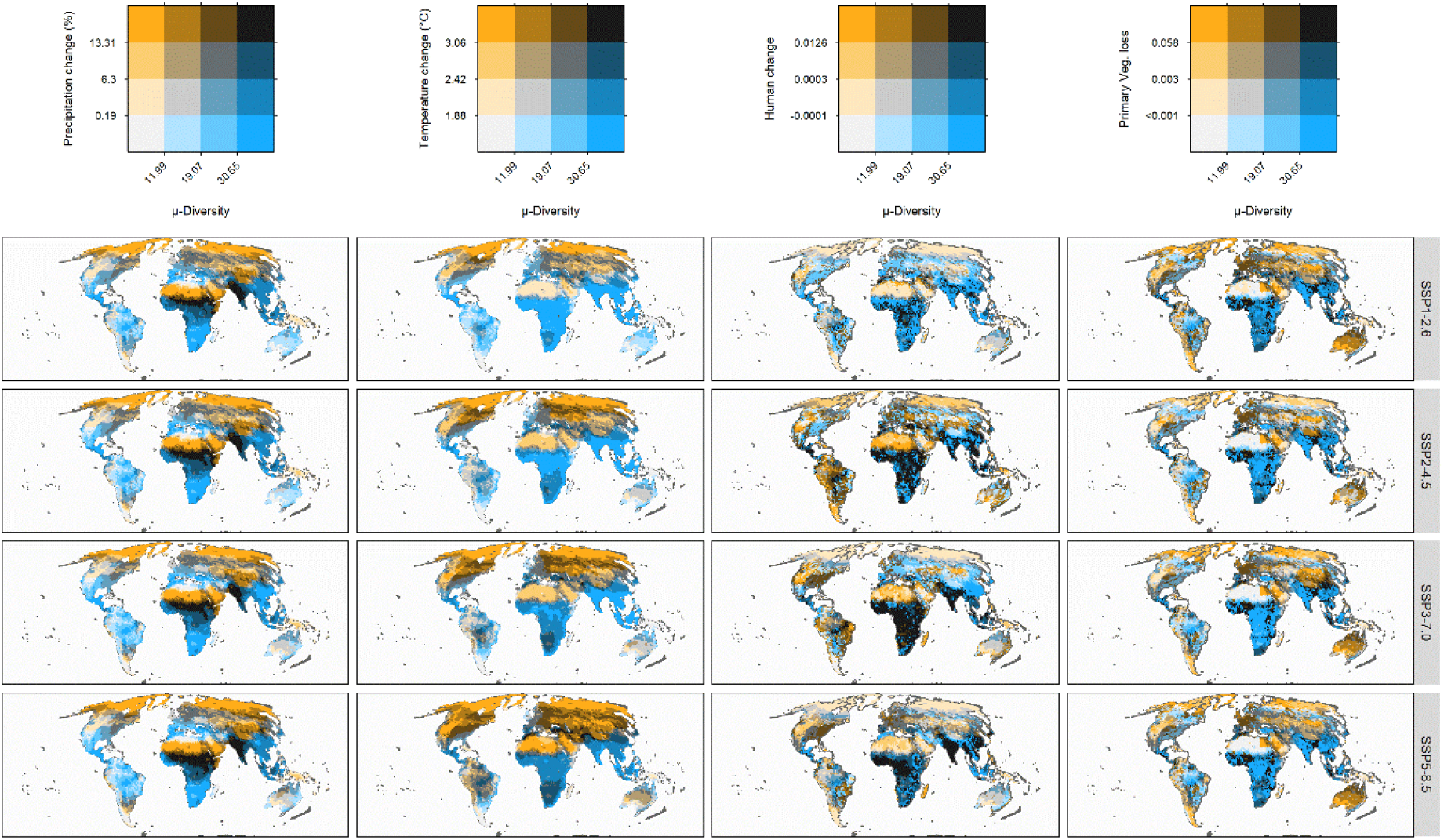
Exposure of mammal μ-Diversity to projected global change across four socioeconomic future scenarios. The y-axis indicates the predicted relative change (i.e. exposure) for each climate and land-use variable, while x-axis represents μ-Diversity. Both axes were expressed in quartiles, darker tones indicate larger exposure in relation to μ-Diversity. We considered four Tier 1 scenarios spanning from a sustainability scenario based on green economy (i.e., “Taking the Green Road”, SSP1-2.6) to a high emission scenario mainly based on fossil fuel exploitation (i.e., “Fossil-fueled Development”, SSP5-8.5).

**Fig. S11.**
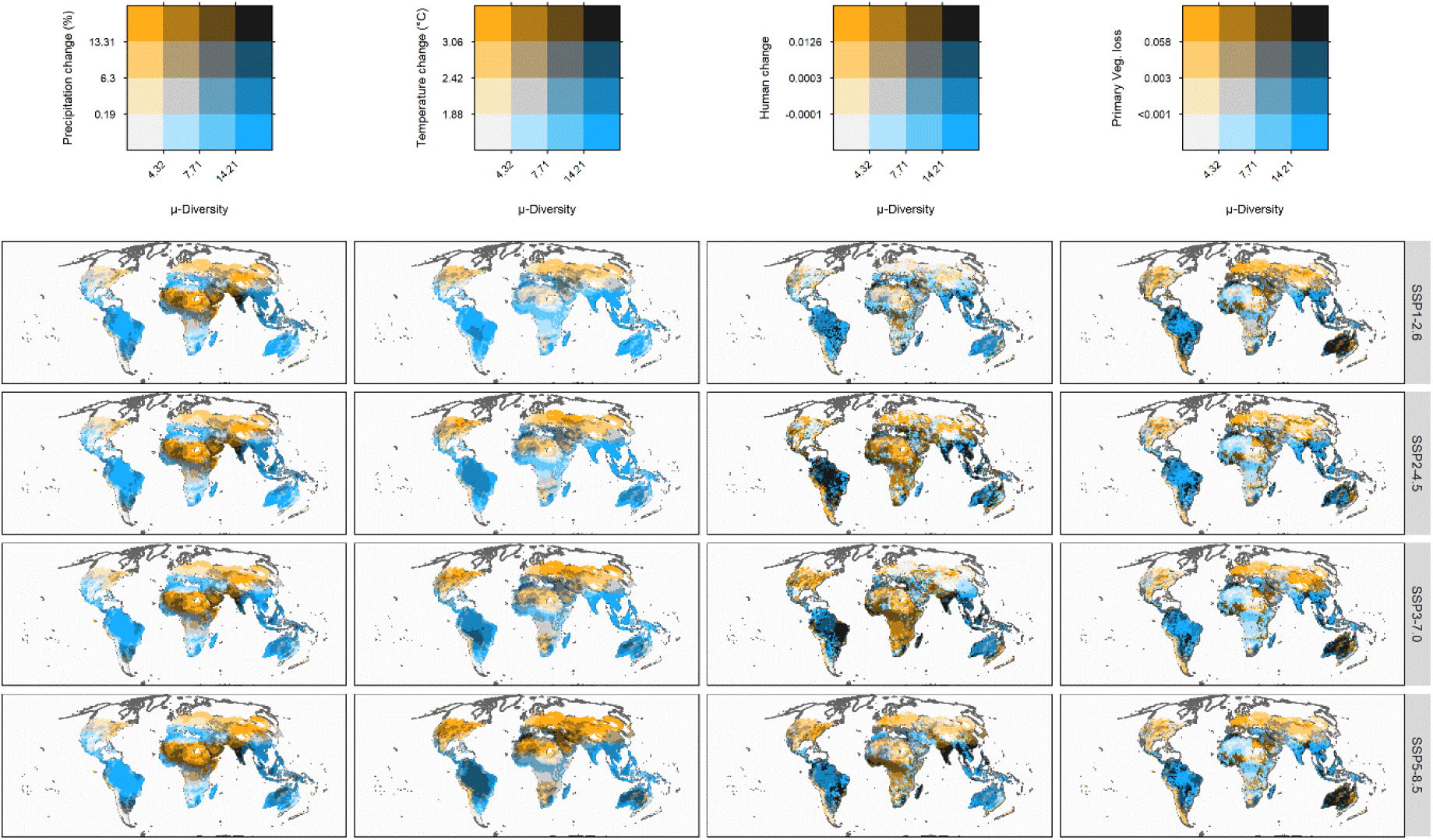
Exposure of reptile μ-Diversity to projected global change across four socioeconomic future scenarios. The y-axis indicates the predicted relative change (i.e. exposure) for each climate and land-use variable, while x-axis represents μ-Diversity. Both axes were expressed in quartiles, darker tones indicate larger exposure in relation to μ-Diversity. We considered four Tier 1 scenarios spanning from a sustainability scenario based on green economy (i.e., “Taking the Green Road”, SSP1-2.6) to a high emission scenario mainly based on fossil fuel exploitation (i.e., “Fossil-fueled Development”, SSP5-8.5).

**Fig. S12.**
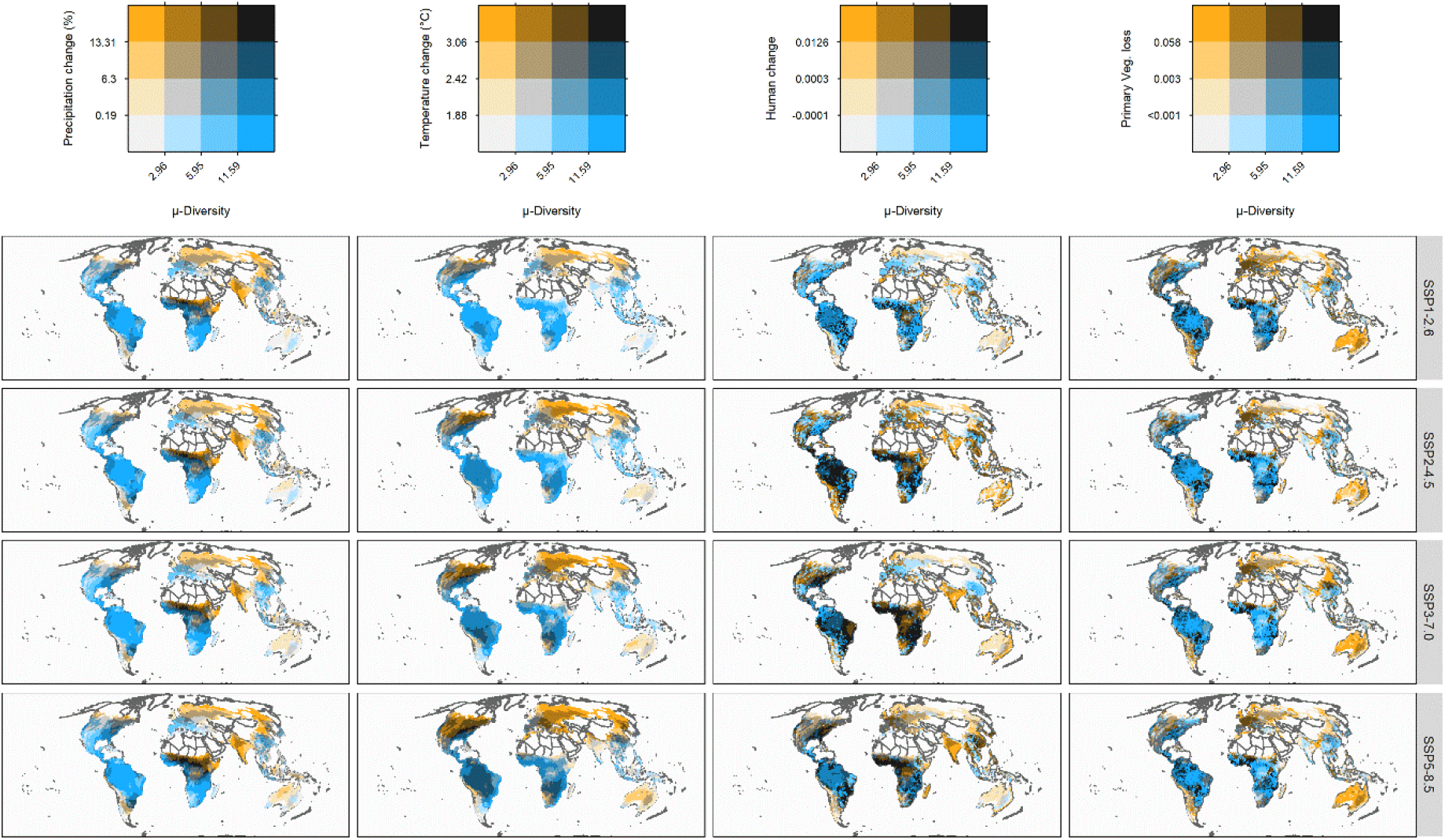
Exposure of amphibian μ-Diversity to projected global change across four socioeconomic future scenarios. The y-axis indicates the predicted relative change (i.e. exposure) for each climate and land-use variable, while x-axis represents μ-Diversity. Both axes were expressed in quartiles, darker tones indicate larger exposure in relation to μ-Diversity. We considered four Tier 1 scenarios spanning from a sustainability scenario based on green economy (i.e., “Taking the Green Road”, SSP1-2.6) to a high emission scenario mainly based on fossil fuel exploitation (i.e., “Fossil-fueled Development”, SSP5-8.5).

**Fig. S13.**
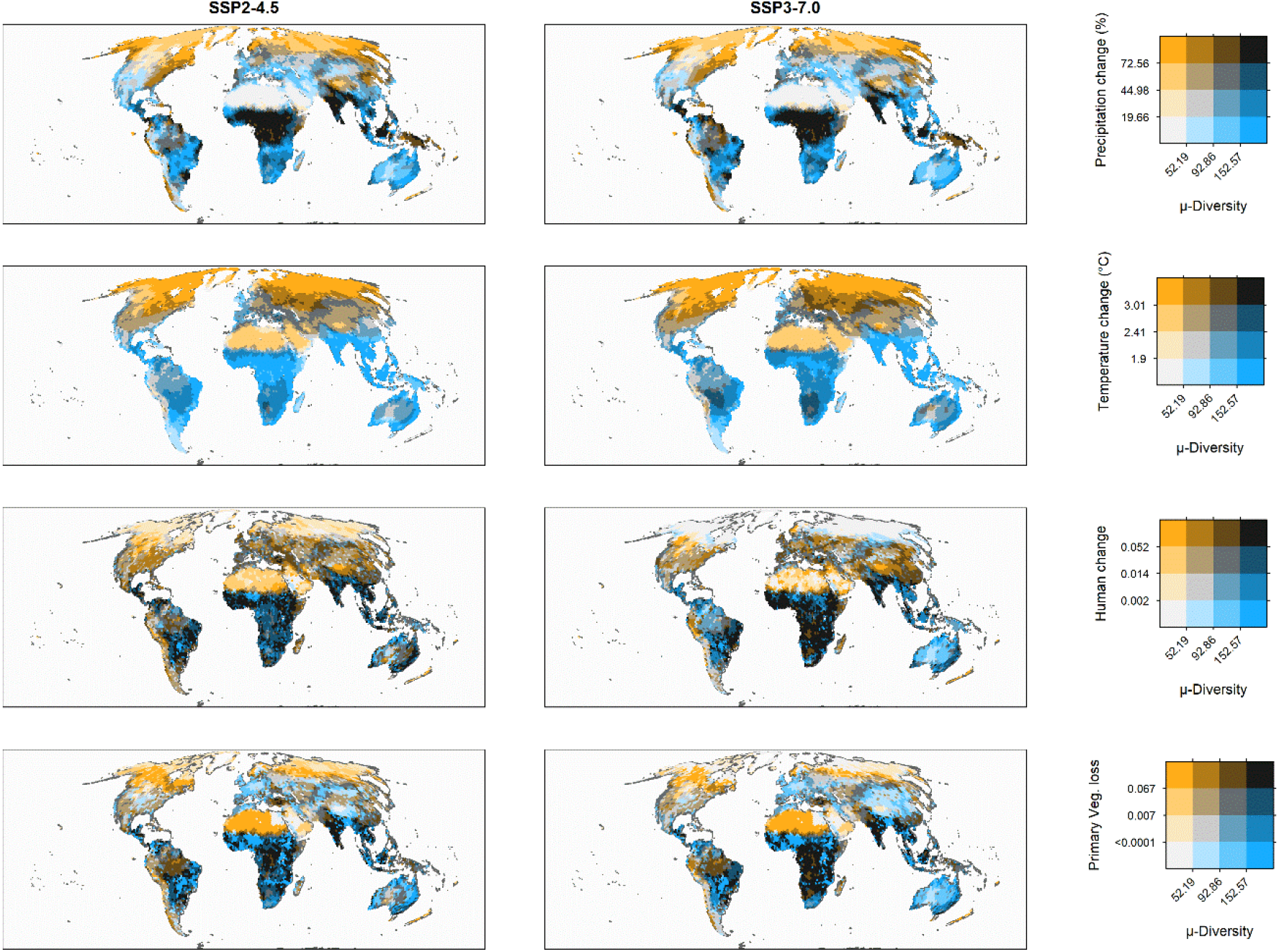
Exposure of tetrapod μ-Diversity to projected global change in the intermediate socioeconomic future scenarios. The y-axis indicates the predicted absolute change (i.e. exposure) for each climate and land-use variable, while x-axis represents μ-Diversity. Both axes were expressed in quartiles, darker tones indicate larger exposure in relation to μ-Diversity. Here are reported only intermediate scenarios, namely SSP2-4.5 and SSP3-7.0, respectively.

**Fig. S14.**
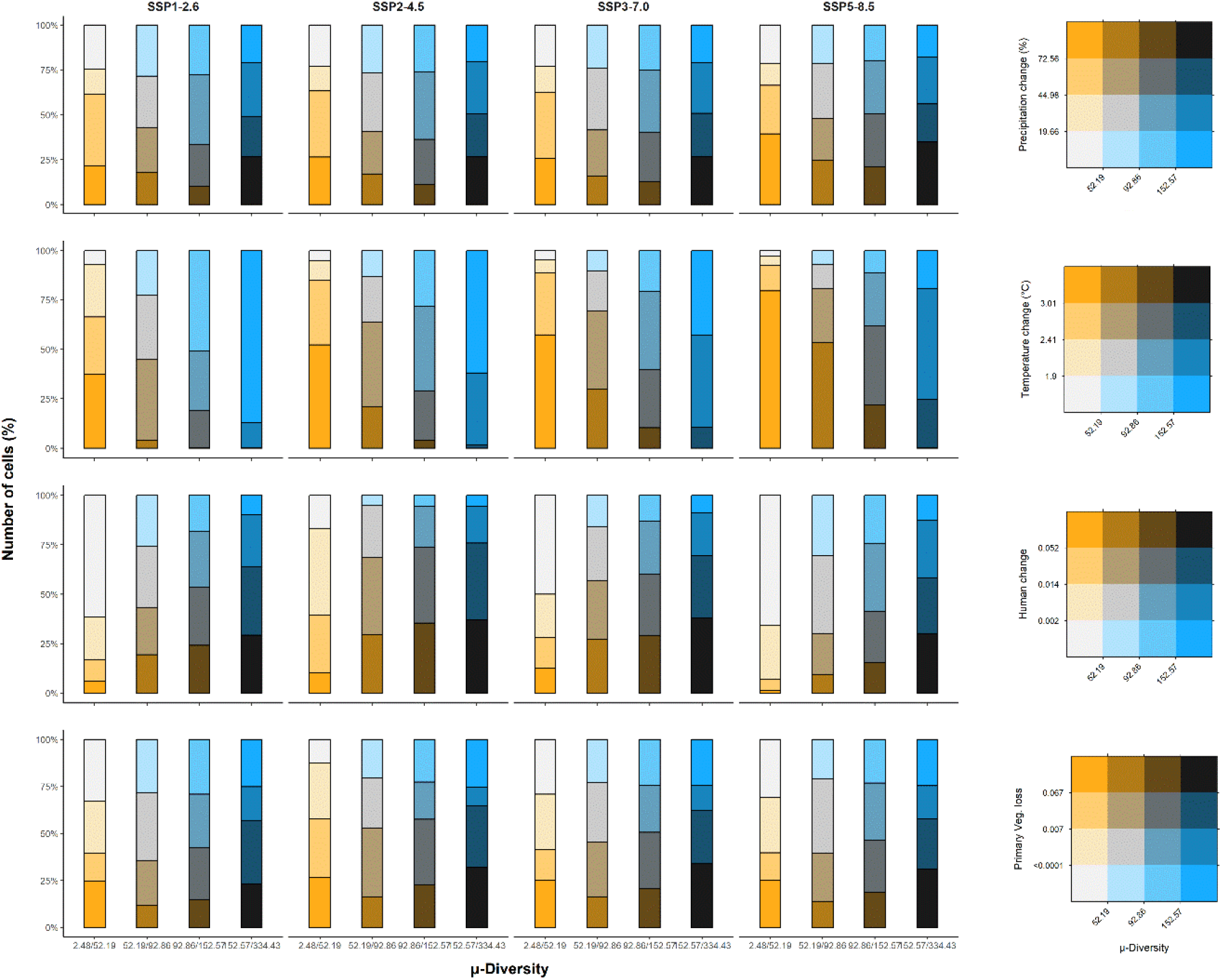
Exposure to projected global change across four socioeconomic future scenarios for each quartile of tetrapod μ-Diversity. The y-axis indicates the proportion of cells (%) belonging to each quartile of the considered variable, while x-axis represents the four quartiles (0–25, 25–50, 50–75, 75–100%) of μ-Diversity. Darker tones indicate larger exposure in relation to the specific quartile of μ-Diversity. We considered four Tier 1 scenarios spanning from a sustainability scenario based on green economy (i.e., “Taking the Green Road”, SSP1-2.6) to a high emission scenario mainly based on fossil fuel exploitation (i.e., “Fossil-fueled Development”, SSP5-8.5).

**Fig. S15.**
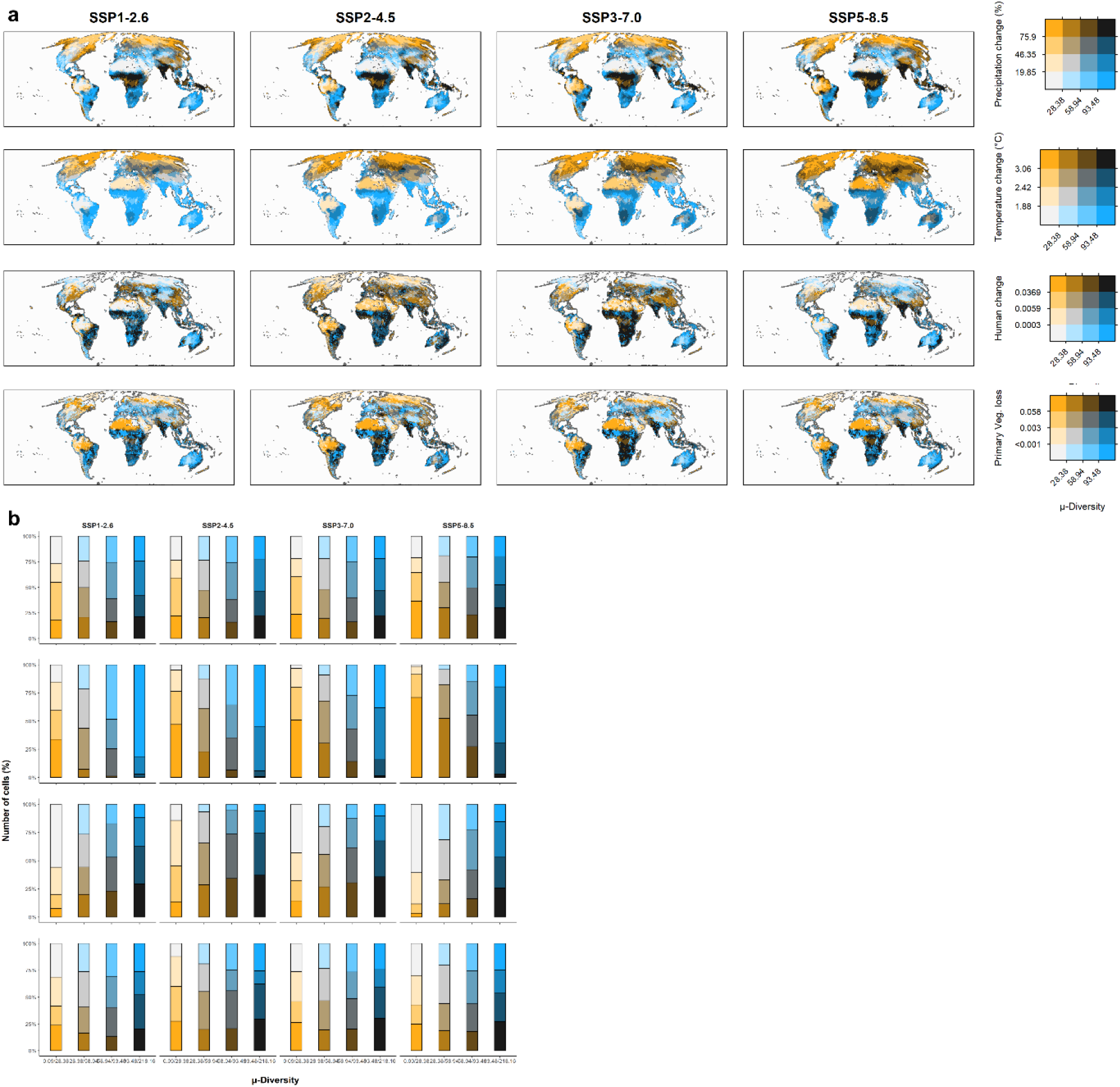
Exposure of avian μ-Diversity to projected global change across four socioeconomic future scenarios. a) The y-axis indicates the predicted absolute change (i.e. exposure) for each climate and land-use variable, while x-axis represents μ-Diversity. Both axes were expressed in quartiles, darker tones indicate larger exposure in relation to μ-Diversity. We considered four Tier 1 scenarios spanning from a sustainability scenario based on green economy (i.e., “Taking the Green Road”, SSP1-2.6) to a high emission scenario mainly based on fossil fuel exploitation (i.e., “Fossil-fueled Development”, SSP5-8.5). b) Relative number of cells (%) belonging to different classes of exposure for each quartile of avian μ-Diversity. The different rows represent the variables reported in panel A following the same order.

**Fig. S16.**
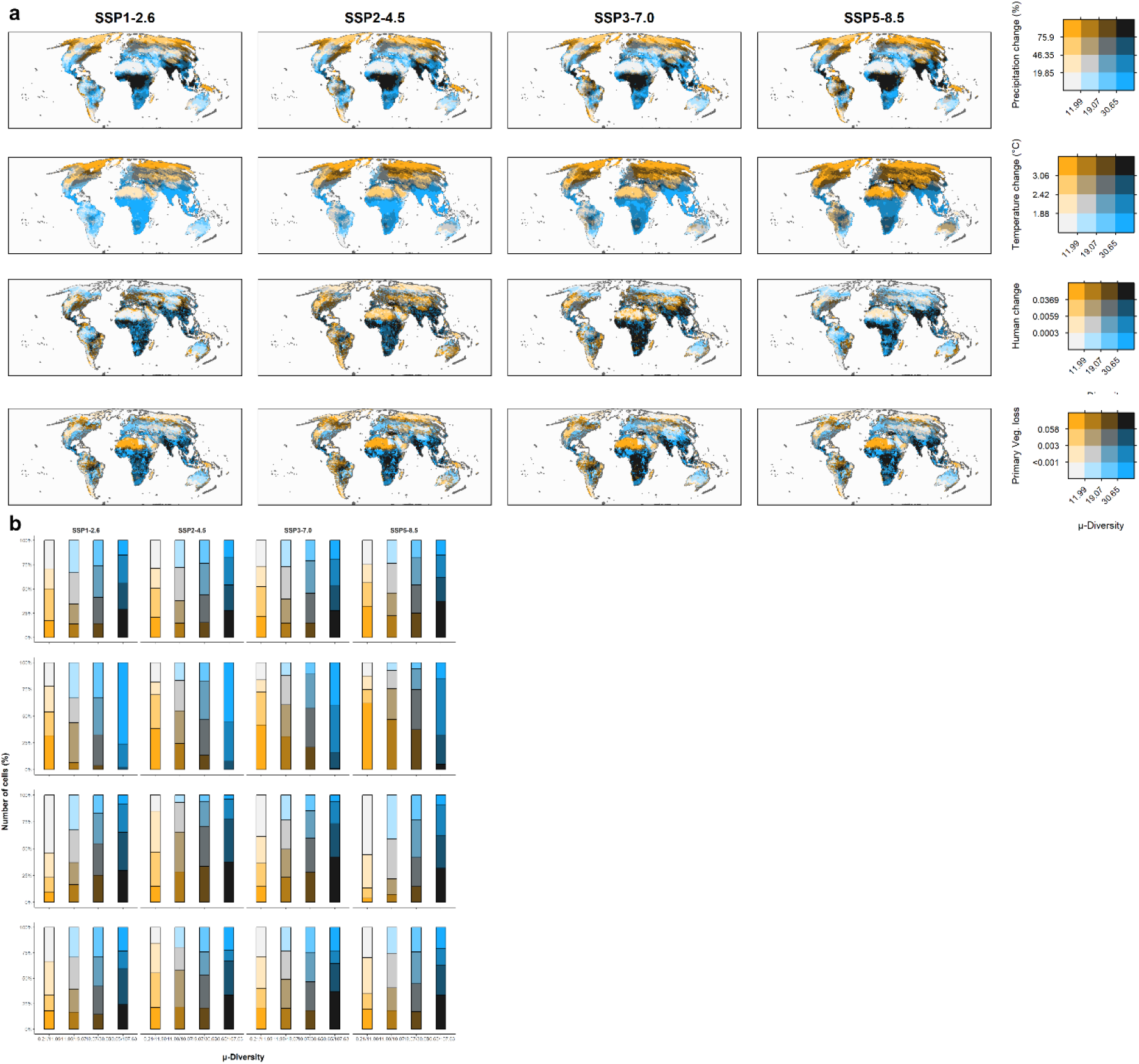
Exposure of mammal μ-Diversity to projected global change across four socioeconomic future scenarios. (A) The y-axis indicates the predicted absolute change (i.e. exposure) for each climate and land-use variable, while x-axis represents μ-Diversity. Both axes were expressed in quartiles, darker tones indicate larger exposure in relation to μ-Diversity. We considered four Tier 1 scenarios spanning from a sustainability scenario based on green economy (i.e., “Taking the Green Road”, SSP1-2.6) to a high emission scenario mainly based on fossil fuel exploitation (i.e., “Fossil-fueled Development”, SSP5-8.5). (B) Relative number of cells (%) belonging to different classes of exposure for each quartile of mammal μ-Diversity. The different rows represent the variables reported in panel A following the same order.

**Fig. S17.**
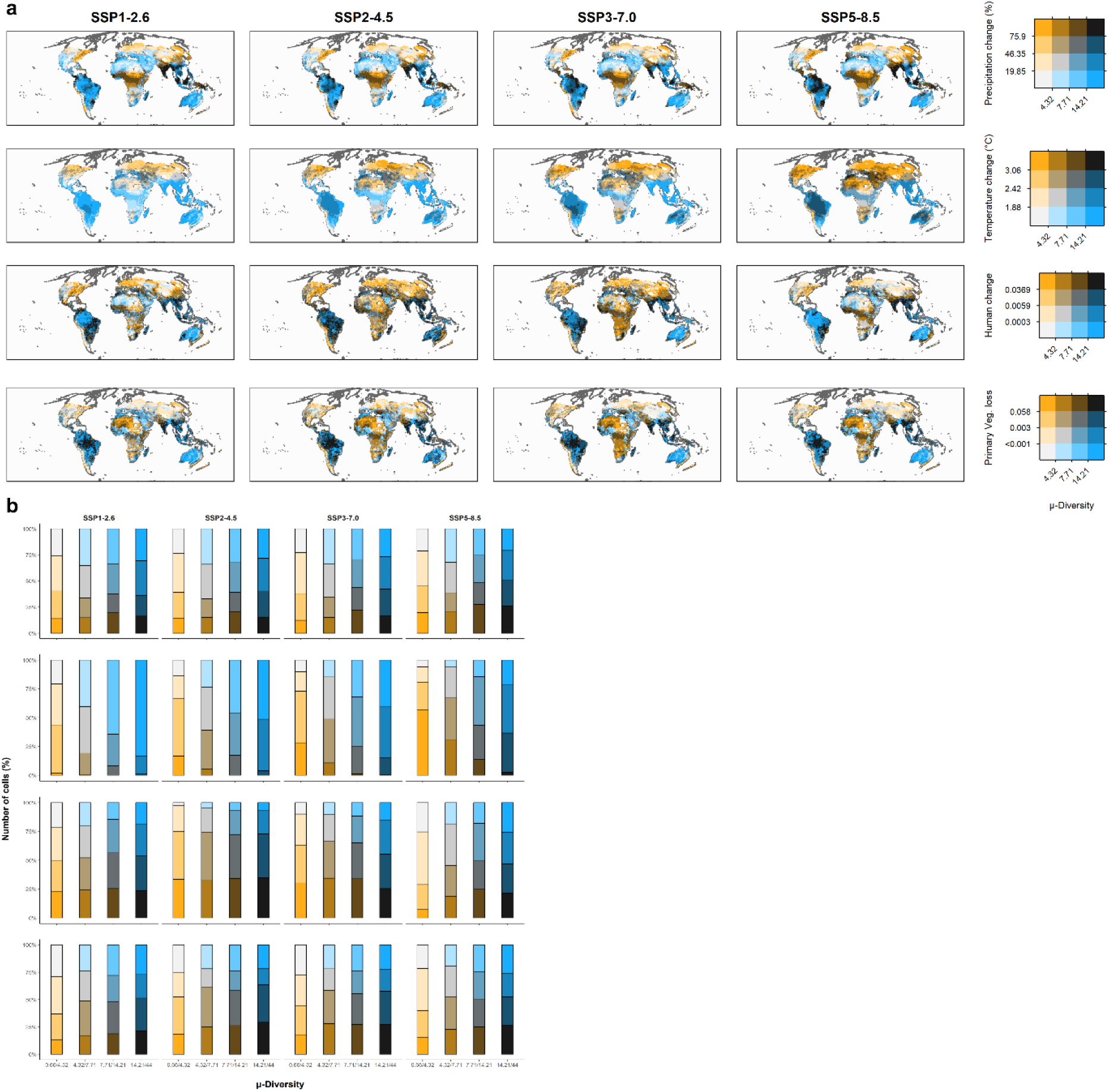
Exposure of reptile μ-Diversity to projected global change across four socioeconomic future scenarios. a) The y-axis indicates the predicted absolute change (i.e. exposure) for each climate and land-use variable, while x-axis represents μ-Diversity. Both axes were expressed in quartiles, darker tones indicate larger exposure in relation to μ-Diversity. We considered four Tier 1 scenarios spanning from a sustainability scenario based on green economy (i.e., “Taking the Green Road”, SSP1-2.6) to a high emission scenario mainly based on fossil fuel exploitation (i.e., “Fossil-fueled Development”, SSP5-8.5). b) Relative number of cells (%) belonging to different classes of exposure for each quartile of reptile μ-Diversity. The different rows represent the variables reported in panel A following the same order.

**Fig. S18.**
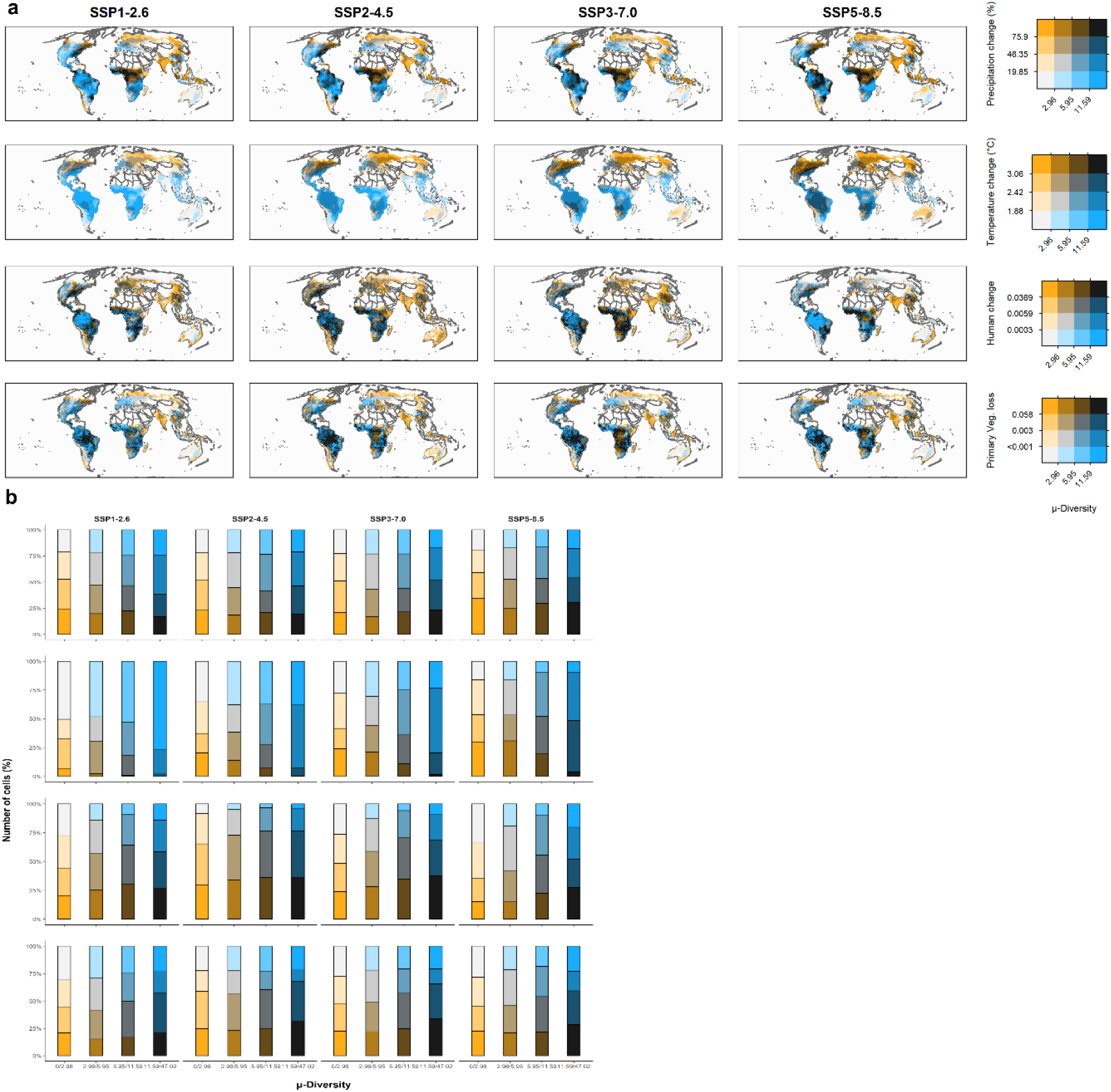
Exposure of amphibian μ-Diversity to projected global change across four socioeconomic future scenarios. (A) The y-axis indicates the predicted absolute change (i.e. exposure) for each climate and land-use variable, while x-axis represents μ-Diversity. Both axes were expressed in quartiles, darker tones indicate larger exposure in relation to μ-Diversity. We considered four Tier 1 scenarios spanning from a sustainability scenario based on green economy (i.e., “Taking the Green Road”, SSP1-2.6) to a high emission scenario mainly based on fossil fuel exploitation (i.e., “Fossil-fueled Development”, SSP5-8.5). (B) Relative number of cells (%) belonging to different classes of exposure for each quartile of amphibian μ-Diversity. The different rows represent the variables reported in panel A following the same order.

**Fig. S19.**
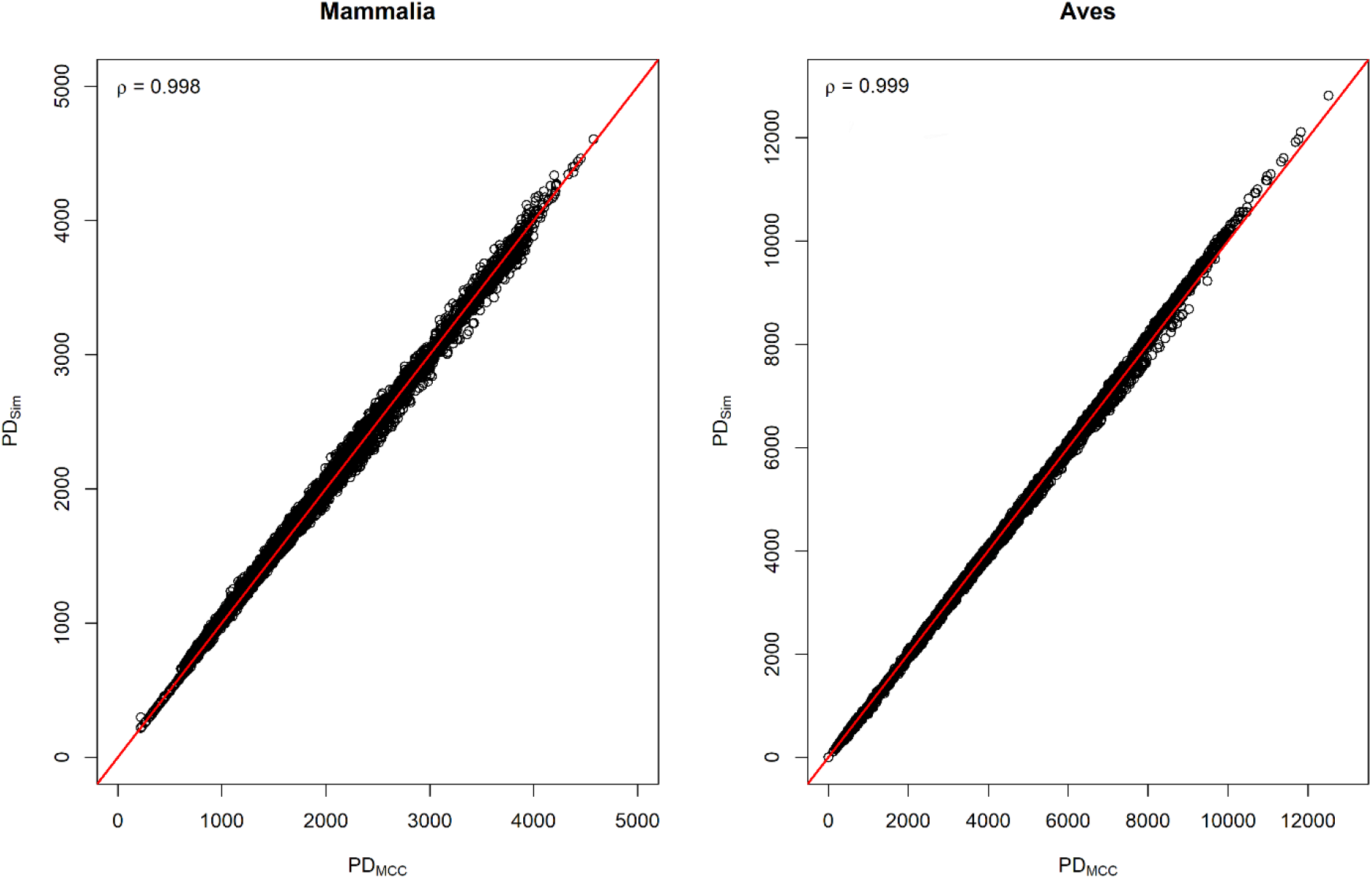
Comparison of phylogenetic diversity values calculated with a maximum clade credibility (PDMCC) tree and PD calculated averaging the values from 100 trees selected from the posterior distribution of mammal and bird phylogenies (PDsim). Red line represents the perfect fit. In both groups, PD values across assemblages were very similar regardless of the method used (Spearman’s ρ > 0.99). We conclude that using a MCC tree should not affect our results.

**Fig. S20.**
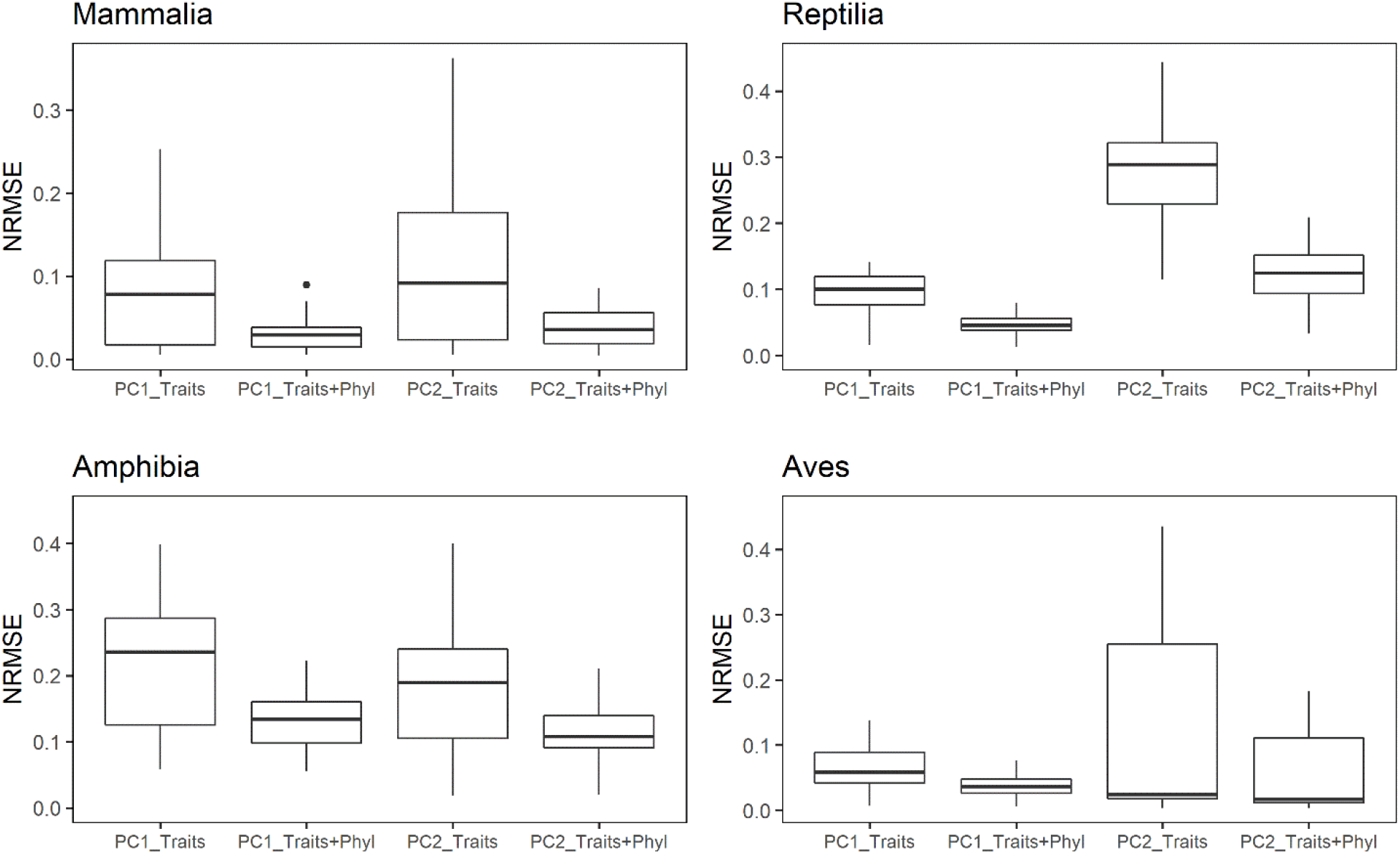
Sensitivity analysis on trait imputation procedure for each taxonomic group. We simulated missing traits (100 repetitions) starting for a subset of species with complete trait data. We then randomly selected 10% of species assigning them the structure of missing values of a random species from the subset of species with missing trait values. Then we combined the three datasets (90% species with complete traits, 10% with simulated NA and the remaining species with non-complete trait information). Here we performed two imputation processes: one based solely on the relationships between functional traits (e.g., PC1_Traits) and another based on the phylogenetic information (e.g., PC1_Traits+Phyl) as described in the methods in the main text. For each dataset obtained, we then computed a functional space using a PCA on which we predicted the position of all species. For only the species with artificial NA, we evaluated the normalized root mean square error (NRMSE) between the original position in the functional space and the position calculated after trait imputation, expressed as the relative range of trait values in the corresponding PC axis.

**Table S1.** Median diversity metric scores for each taxonomic group and for all tetrapods and the relative coverage in terms of number of species.

| Clade | Species with functional,<br>phylogenetic and range<br>data | Total species | % total species<br>included in this study |
| --- | --- | --- | --- |
| **Tetrapoda** | **17,341** | **35,283** | **49%** |
| Mammalia | 3,912 | ~5,692 | 69% |
| Amphibia | 3,239 | 7,776 | 42% |
| Reptilia | 3,338 | 10,845 | 31% |
| Aves | 6,852 | 10,970 | 62% |

**Table S2.** Pearson’s correlations between diversity metrics for each taxonomic group. All the correlations were spatially corrected. TD refers to species richness, sesPD to the standardized effect size (SES) related to phylogenetic diversity (PD) while sesFD to the effect size related to functional richness (FD). SES were computed as follows: SES = (Metric_obs_−mean(Metric_null_))/ SD(Metric_nul_l)_l_.

| Taxonomic group | TD | sesPD | sesFD |  |
| --- | --- | --- | --- | --- |
| *Mammalia* | - |  |  | TD |
|  | -0.69*** | - |  | sesPD |
|  | 0.12 | 0.16 | - | sesFD |
| *Amphibia* | - |  |  | TD |
|  | -0.75*** | - |  | sesPD |
|  | 0.22 | 0.01 | - | sesFD |
| *Reptilia* | - |  |  | TD |
|  | -0.88** | - |  | sesPD |
|  | -0.22 | 0.39 | - | sesFD |
| *Aves* | - |  |  | TD |
|  | -0.33 | - |  | sesPD |
|  | -0.52*** | 0.30* | - | sesFD |

\*\*\* = $P < 0.001$ ; \*\* = $P < 0.01$ ; \* = $P < 0.05$

**Table S3.** Model output showing the variable importance expressed using Root Mean Squared Error loss (RMSE, median ± SD) for each predictor considering tetrapod pooled and each taxonomic group independently for all three diversity facets. NRMSE and R^2^ were obtained using spatial cross-validation, NRMSE is the Normalized Root Mean Squared Error normalized over the range of the response variable in each dataset. *N* represents the number of grid cells used to train the models. Please note that for all the five datasets, only cells without missing values were used.

|  |  | **Tetrapoda** | **Mammalia** | **Amphibia** | **Reptilia** | **Aves** |
| --- | --- | --- | --- | --- | --- | --- |
| *μ-Diversity* | Biome variation | 28.891 ± 0.193 | 11.804 ± 0.063 | 4.798 ± 0.034 | 4.881 ± 0.02 | 20.170 ± 0.102 |
| | Climate variation | 40.036 ± 0.31 | 14.363 ± 0.095 | 5.230 ± 0.034 | 5.509 ± 0.034 | 31.587 ± 0.253 |
| | Agricultural onset | 31.986 ± 0.183 | 13.187 ± 0.081 | 5.253 ± 0.033 | 5.229 ± 0.023 | 25.853 ± 0.191 |
| | HFP | 31.692 ± 0.22 | 13.249 ± 0.101 | 5.370 ± 0.04 | 4.855 ± 0.016 | 24.557 ± 0.192 |
| | PET | 38.818 ± 0.33 | 13.313 ± 0.09 | 5.210 ± 0.029 | 5.732 ± 0.04 | 29.09 ± 0.219 |
| | Soil humidity | 31.223 ± 0.192 | 12.827 ± 0.074 | 5.250 ± 0.035 | 5.278 ± 0.025 | 21.972 ± 0.108 |
| | Forest cover | 30.895 ± 0.226 | 12.129 ± 0.075 | 4.929 ± 0.027 | 5.134 ± 0.023 | 22.303 ± 0.118 |
| | **$R^2$** | 0.80 ± 0.03 | 0.69 ± 0.05 | 0.76 ± 0.08 | 0.69 ± 0.05 | 0.73 ± 0.03 |
| | **NRMSE** | 0.14 ± 0.03 | 0.19 ± 0.06 | 0.14 ± 0.10 | 0.21 ± 0.07 | 0.15 ± 0.04 |
|  | **$N$** | 5,039 | 5,001 | 3,498 | 4,205 | 5,027 |
| *Taxonomic Diversity* | Biome variation | 74.684 ± 0.668 | 20.327 ± 0.113 | 10.415 ± 0.087 | 15.313 ± 0.087 | 47.988 ± 0.333 |
| | Climate variation | 107.847 ± 1.011 | 26.568 ± 0.2 | 12.191 ± 0.108 | 16.768 ± 0.108 | 68.424 ± 0.611 |
| | Agricultural onset | 80.036 ± 0.506 | 21.848 ± 0.133 | 11.306 ± 0.091 | 17.128 ± 0.099 | 51.441 ± 0.276 |
| | HFP | 82.411 ± 0.686 | 20.644 ± 0.105 | 11.426 ± 0.09 | 16.097 ± 0.105 | 52.249 ± 0.377 |
| | PET | 102.538 ± 0.844 | 24.685 ± 0.17 | 11.840 ± 0.078 | 17.074 ± 0.124 | 57.900 ± 0.436 |
| | Soil humidity | 94.426 ± 0.767 | 24.846 ± 0.192 | 11.325 ± 0.08 | 15.814 ± 0.068 | 60.987 ± 0.467 |
| | Forest cover | 100.270 ± 1.133 | 25.238 ± 0.21 | 11.654 ± 0.087 | 16.626 ± 0.098 | 60.806 ± 0.584 |
| | **$R^2$** | 0.85 ± 0.02 | 0.80 ± 0.04 | 0.79 ± 0.05 | 0.79 ± 0.06 | 0.82 ± 0.03 |
| | **NRMSE** | 0.10 ± 0.07 | 0.12 ± 0.06 | 0.10 ± 0.07 | 0.22 ± 0.10 | 0.11 ± 0.06 |
|  | **$N$** | 5,039 | 5,001 | 3,498 | 4,205 | 5,027 |
| *Ecological Distinctiveness* | Biome variation | 0.074 ± 0 | 0.136 ± 0 | 0.134 ± 0 | 0.147 ± 0.001 | 0.123 ± 0.001 |
| | Climate variation | 0.081 ± 0 | 0.150 ± 0.001 | 0.162 ± 0.001 | 0.168 ± 0.001 | 0.131 ± 0.001 |
| | Agricultural onset | 0.083 ± 0 | 0.158 ± 0.001 | 0.146 ± 0.001 | 0.171 ± 0.001 | 0.131 ± 0.001 |
| | HFP | 0.076 ± 0 | 0.155 ± 0.001 | 0.136 ± 0 | 0.151 ± 0.001 | 0.125 ± 0.001 |
| | PET | 0.088 ± 0 | 0.160 ± 0.001 | 0.141 ± 0.001 | 0.168 ± 0.001 | 0.154 ± 0.001 |
| | Soil humidity | 0.078 ± 0 | 0.137 ± 0 | 0.134 ± 0 | 0.152 ± 0.001 | 0.141 ± 0.001 |
| | Forest cover | 0.086 ± 0.001 | 0.141 ± 0.001 | 0.143 ± 0.001 | 0.159 ± 0.001 | 0.143 ± 0.001 |
| | **$R^2$** | 0.74 ± 0.05 | 0.63 ± 0.03 | 0.56 ± 0.03 | 0.65 ± 0.06 | 0.72 ± 0.03 |
| | **NRMSE** | 0.18 ± 0.03 | 0.23 ± 0.05 | 0.22 ± 0.05 | 0.28 ± 0.05 | 0.20 ± 0.04 |
|  | **$N$** | 5,039 | 5,001 | 3,498 | 4,205 | 5,027 |

**Table S4.** Pearson’s correlation between μ-Diversity and the project global change. Correlations were calculated for all tetrapods and for each taxonomic group between μ-Diversity and the future climatic and land-use absolute changes; positive correlations suggest a higher exposure at higher levels of diversity and viceversa. All the correlations were spatially corrected.

| Scenario | Variable | Tetrapoda | Aves | Mammalia | Reptilia | Amphibia |
| --- | --- | --- | --- | --- | --- | --- |
| SSP1-2.6 | Temperature change | -0.65*** | -0.57*** | -0.46*** | -0.47** | -0.19 |
|  | Precipitation change | 0.09 | -0.04 | 0.22** | 0.04 | -0.1 |
|  | Human change | 0.31*** | 0.28*** | 0.25*** | 0.00 | 0 |
|  | Primary veg. loss | 0.00 | -0.02 | 0.10 | 0.02 | -0.07 |
| SSP2-4.5 | Temperature change | -0.65*** | -0.57*** | -0.47*** | -0.46** | -0.17 |
|  | Precipitation change | 0.08 | 0.04 | 0.18* | 0.04 | -0.02 |
|  | Human change | 0.34*** | 0.33*** | 0.24*** | 0.03 | 0 |
|  | Primary veg. loss | 0.12 | 0.07 | 0.19** | 0.05 | 0 |
| SSP3-7.0 | Temperature change | -0.64*** | -0.57*** | -0.48*** | -0.45** | -0.14 |
|  | Precipitation change | 0.09 | 0.05 | 0.15* | 0.06 | 0.06 |
|  | Human change | 0.39** | 0.32** | 0.44*** | -0.11 | 0.13 |
|  | Primary veg. loss | 0.23* | 0.16 | 0.33** | -0.04 | 0.08 |
| SSP5-8.5 | Temperature change | -0.65*** | -0.58*** | -0.47*** | -0.44* | -0.16 |
|  | Precipitation change | 0.12 | 0.06 | 0.20* | 0.09 | 0 |
|  | Human change | 0.39*** | 0.29*** | 0.44*** | 0.10 | 0.07 |
|  | Primary veg. loss | 0.18* | 0.11* | 0.30*** | 0.06 | 0 |

\* = $P < 0.05$ ; \*\* = $P < 0.01$ ; \*\*\* = $P < 0.001$

**Table S5.** Correlation matrix (Spearman’s ρ) of the predictors used in random forest models.

|  | Climate<br>variation | Biome<br>variation | PET | Soil<br>humidity | HFP | Forest<br>cover | Agricultural<br>onset |
| --- | --- | --- | --- | --- | --- | --- | --- |
| Climate variation | - |  |  |  |  |  |  |
| Biome variation | -0.4 | - |  |  |  |  |  |
| PET | -0.6 | 0.2 | - |  |  |  |  |
| Soil humidity | -0.2 | 0.3 | -0.3 | - |  |  |  |
| HFP | -0.2 | 0.2 | 0.2 | 0.2 | - |  |  |
| Forest cover | -0.1 | 0.3 | -0.3 | 0.7 | 0.0 | - |  |
| Agricultural onset | -0.1 | 0.0 | -0.1 | 0.2 | -0.4 | 0.3 | - |

**Table S6.** List of traits used in this study along with their completeness.

| Taxonomic group | Trait | Abbreviations [unit measure] | Number of species | Completeness (%) |
| --- | --- | --- | --- | --- |
| Mammalia | Adult body mass | bm [g] | 4,651 | 93.90 |
|  | Longevity | long [years] | 2,614 | 52.78 |
|  | Number of offspring per litter | ls [n] | 3,511 | 70.89 |
|  | Litter per year | ly [n] | 2,146 | 43.44 |
|  | Weaning length | wea[days] | 2,043 | 41.25 |
|  | Time to female maturity | fmat [days] | 2,000 | 40.38 |
|  | Snout-vent length (distance from the tip of the snout to the tail base) | svl [cm] | 3,921 | 79.16 |
|  | Gestation length | gest [days] | 2,220 | 44.82 |
| Amphibia | Maximum litter size | ls [n] | 1,623 | 23.95 |
|  | Offspring size | os [mm] | 1,330 | 19.63 |
|  | Age at maturity | am [years] | 384 | 5.67 |
|  | Body size | bs [mm] | 5,227 | 77.14 |
| Reptilia | Longevity | long [years] | 1,214 | 18.49 |
|  | Number of clutches per year | ly [n] | 1,018 | 15.5 |
|  | Clutch size | ls [n] | 2,675 | 40.73 |
|  | Adult body mass | bm [g] | 2,494 | 37.98 |
|  | Incubation time | inc [days] | 1,369 | 20.85 |
|  | Snout-vent length (distance from the tip of the snout to the opening of the cloaca) | svl [cm] | 5,140 | 78.27 |
| Aves | Adult body mass | bm [g] | 9,532 | 97.25 |
|  | Number of clutches per year | ly [n] | 1,784 | 18.20 |
|  | Clutch size | ls [n] | 6,892 | 70.31 |
|  | Egg mass | em [g] | 4,888 | 49.87 |
|  | Fledging age | fa [days] | 1,844 | 18.81 |
|  | Snout-vent length (distance from the tip of the snout to the opening of the cloaca) | svl [cm] | 1,615 | 16.48 |
|  | Longevity | long [years] | 1,672 | 17.06 |
|  | Incubation time | inc [days] | 2,269 | 23.15 |

